# Independent Component Analysis Outperforms Seed-Based Approach in Detecting fNIRS-based Resting-State Functional Connectivity

**DOI:** 10.64898/2026.02.18.705990

**Authors:** Foivos Kotsogiannis, Sophie Raible, João Pereira, Armin Heinecke, Simona Klinkhammer, Bettina Sorger, Michael Lührs

## Abstract

**Significance:** Resting-state functional connectivity (RSFC) is an important measure in advancing our understanding of brain function and development as well as various neurological and mental disorders. Studying RSFC with functional near-infrared spectroscopy (fNIRS) offers several advantages over functional magnetic resonance imaging (fMRI), especially for clinical and pediatric populations. However, the optimal strategy to estimate RSFC based on fNIRS, particularly in identifying reliable connectivity patterns across chromophores, remains unclear. Establishing robust analysis approaches is essential for reliable and clinically meaningful applications.

**Aim:** This study systematically evaluated commonly used analysis methods regarding their effectiveness to detect RSFC patterns within the motor network using both oxygenated (HbO) and deoxygenated (HbR) hemoglobin signals.

**Approach:** Near whole-head resting-state fNIRS data were analyzed from 38 participants. RSFC was estimated with five analytical approaches: three seed-based methods (SBA-GLM, SBA-GLM with respiratory regression, and SBA-correlation) and two independent component analyses (ICA) approaches using two different contrast functions. Performance was assessed via receiver operating characteristic analyses based on both anatomical and functional definitions of motor-related connectivity. Areas under the curves (AUC) were statistically compared with DeLong’s test, and the spatial similarity between HbO and HbR RSFC was quantified by correlating RSFC patterns from the two chromophores.

**Results:** Across reference definitions and chromophores, ICA consistently achieved higher performance (AUC = 0.82–0.96) in detecting motor-related RSFC than SBA (AUC = 0.63–0.86). Significant differences emerged when functionally defined connectivity references were used, with ICA outperforming SBA across chromophores. Under certain condition, correlational-SBA (AUC = 0.66–0.86) significantly outperformed GLM-based methods (AUC = 0.63–0.85). Finally, ICA results demonstrated greater spatial similarity between obtained HbO and HbR RSFC patterns (*r* = 0.90–0.92) than SBA (*r* = 0.84–0.86), indicating higher cross-chromophore consistency.

**Conclusions:** ICA provides a robust and consistent framework for estimating fNIRS-based RSFC across both HbO and HbR, outperforming SBA in accuracy and cross-chromophore consistency. While correlational-SBA offers a computationally efficient alternative and outperforms GLM-based methods, ICA should be preferred when reliable and chromophore-consistent RSFC estimates are required. Importantly, these findings demonstrate that HbR contains RSFC information comparable to HbO and highlights the critical role of analytical strategy and reference definition in RSFC evaluation. Collectively, these results contribute to the methodological standardization of fNIRS-based RSFC and support its use in future neuroscientific and clinical applications.

## 1 Introduction

The brain is a highly interconnected system, where anatomically separated areas are temporally dependent in terms of neural activity, a phenomenon known as functional connectivity (FC),^1–4^ which allows the continuous and efficient flow of information.^3–5^ Research on FC could improve our understanding of brain function and development^3,6,7^ and assist in the diagnosis and treatment of mental and neurological disorders.^8–10^

With methodological advances, our ability to identify functionally interconnected networks has improved. With resting-state (RS) paradigms, participants are not engaging in any task, which enables the detection of FC between distinct regions based on their synchronous low frequency oscillations (∼0.01–0.1 Hz).^4,6^ The feasibility and simplicity of this technique increase replicability of findings^6,8^ while minimizing the effort required from participants, which is advantageous for studying clinical^11^ and pediatric populations.^6,7,11^

Resting-state functional connectivity (RSFC) has been extensively studied with fMRI.^1,5,10,11^ Recently, functional near-infrared spectroscopy (fNIRS) has emerged as a valid and reliable alternative with distinct advantages over fMRI,^3,5,6,12–17^ making fNIRS particularly well-suited for clinical populations^13,18,19^ and, in effect, for studying FC.

fNIRS relies on the hemodynamic response and measures relative concentrations of oxygenated (HbO) and deoxygenated (HbR) hemoglobin with near-infrared (NIR) light. Source optodes emit NIR light, which penetrates through scalp and bone to reach the cerebral cortex.^4,6,13,19^ As light travels it is either absorbed or scattered by tissue.^4,13,19^ With appropriate source-detector spacing and optimal wavelengths in the 650-950 nm range, NIR light can penetrate to the cerebral cortex, where HbO and HbR absorb it differently. Exploiting these distinct absorption spectra, the absorption of NIR radiation by HbO and HbR can be differentiated and quantified^4,13^ by capturing the returning light via detectors^4,6,13,19^ and comparing its intensity to the initially emitted light.^4^

Since fNIRS is a mobile technology, more ecologically valid studies can be conducted as compared to fMRI.^13^ Participants can move freely in the natural environments, the system is considerably more tolerant to motion artifacts, it is silent, more accessible due to lower cost,^13,18,19^ and easier to use.^13^ It can be utilized across diverse populations, including those with medical conditions (e.g., neurological and mental disorders), infants, and those unsuitable for MRI (e.g., with metal implants, claustrophobia, etc.).^13,18^ Lastly, fNIRS has a higher sample rate compared to fMRI, but being restricted to outer layers of the cortex, it has lower spatial resolution and brain coverage.^13,18,19^

### 1.1 RSFC with fNIRS

Various techniques have been used to estimate RSFC with fNIRS, with the most widely used methods being seed-based analysis (SBA), independent component analysis (ICA), and graph theory analysis.^3,4,20^ Other methods such as artificial neural networks (ANN), convolutional neural networks (CNN),^21^ and Bayesian networks analysis^22^ show potential in determining RSFC.

Despite growing interest in RSFC research using fNIRS, the optimal analytical strategy remains unclear, especially in terms of identifying reliable patterns across chromophores. The current study will focus on addressing this gap by systematically comparing commonly used methods (i.e., SBA and ICA) to estimate RSFC from both HbO and HbR signals.

Beyond methodological advancements, the current work may offer translational value for clinical applications. Currently, fMRI-based functional connectivity analysis is primarily used for presurgical planning, and its broader diagnostic and prognostic potential remain underutilized.^11,23^

This is partly driven by limitations inherent to fMRI (e.g., susceptibility to motion artifacts, operational costs, limited scanner availability, etc.) and the complexity of data analysis.^24^ As discussed earlier, fNIRS is less affected by these constraints and may prove a valuable alternative in assessing FC. Establishing a systematic evaluation and potential standardization of RSFC estimation could therefore facilitate the integration of fNIRS-RSFC in clinical practice, increasing its utility in psychiatric and neurological assessment and rehabilitation.

### 1.2 SBA

SBA estimates the functional relationship between a predefined seed and all other measured locations (voxels in fMRI, channels in fNIRS) across the brain.^3^ A stronger connection between the seed and another voxel/channel indicates a higher connectivity between their corresponding brain regions. Many studies have investigated the feasibility of SBA to produce reliable RSFC maps.

White et al. (2009) were among the first to apply this method in fNIRS, using SBA to evaluate RSFC and validate this approach (*N* = 5).^14^ Based on functional-localization procedures, channels in the motor and visual regions (identified as the most-activated fNIRS channels for each functional task) were defined as the seeds. After regressing the global signal, the time-course of the seed was correlated with all other fNIRS channels to estimate connectivity, which yielded similar RSFC patterns to the fMRI literature at the time. Similarly, Mesquita et al. 2010 used SBA to detect the RSFC patterns in frontal, sensorimotor, and visual regions (*N* = 9; all male).^12^ By visually selecting seed channels for each region, the authors showed that fNIRS can detect FC within regions and also capture inter-regional connectivity.

In parallel with these early validation efforts, Lu et al. (2010) estimated the RSFC of sensorimotor and auditory areas with SBA, but this time using the general linear model (GLM; *N* = 29).^17^ As in previous studies, a functional-localization procedure was used to define the seeds for each task, and the seeds’ time course was used as a predictor in the GLM. With this approach, the authors were able to determine the RSFC patterns of the sensorimotor and auditory cortex successfully. Their paradigm was later replicated^15,21^ and tested for robustness, showing high reliability estimates for RSFC patterns in high-order regions.^16^ These studies provide evidence supporting the suitability of SBA in fNIRS to detect RSFC patterns in lower- and higher-order regions, which is further supported by simultaneous fNIRS-fMRI recordings using similar methods.^20^

Nonetheless, methodological inconsistencies exist in SBA, such as the statistical approach used to determine RSFC (i.e., GLM *vs.* Pearson’s *r*). Moreover, most prior studies were not conducted on large representative samples. In addition, some only analyzed HbO because of its high signal-to-noise ratio (SNR),^16,21^ hindering our understanding of HbR’s role in FC, which might hold important information regarding brain activity.^22^ Other limitations involved not regressing out confounding signals recorded from superficial scalp layers and the complete removal of global signal^12,14^ despite current evidence that the global signal contains meaningful information about RSFC.^4^

### 1.3 ICA

While SBA is a reliable approach to identify RSFC, it ignores the interaction of multiple regions because it relies on a predetermined seed and its relationship to the neighboring channels.^25^ An alternative method is ICA, which is designed to extract information from mixed and unknown sources, reduce noise, and identify functionally interconnected networks.^3^ ICA separates true RS signals from random components. It can identify interactions of functional networks and does not necessarily require a functional-localization procedure, making it more convenient.^11^

H. Zhang et al. (2010) demonstrated ICA’s feasibility and advantage over SBA with GLM in detecting RSFC in sensorimotor and visual regions with fNIRS (*N* = 16).^15^ The authors visually identified the IC expressing increased RSFC (i.e., showing low-frequency activity in bilateral motor area) and assessed the performance of ICA and SBA in generating RSFC maps with receiver operating characteristic (ROC) curves. Based on a structural MRI, the sensorimotor and visual regions were selected as golden standards. When inspecting the RSFC from the HbO signal, ICA had higher sensitivity and specificity than SBA, a finding that was successfully replicated by Behboodi et al. (2019) (*N* = 10; for HbO only).^21^ Further, H. Zhang et al. (2010) compared the similarity of RSFC patterns across chromophores, assuming a strong linear relationship between the two should exist given that FC derives from the same region.^15^ Compared to SBA, ICA yielded higher similarity between HbO/HbR RSFC patterns, suggesting that it is more reliable in RSFC estimation. Although ICA appears more promising than SBA, it remains unclear if noise reduction with short channels could improve SBA to match ICA. Additionally, ICA is time-consuming, especially with large samples^21^ and may remove true signal,^26,27^ suggesting the need for alternative approaches.

### 1.4 Present Study

While the performance of SBA-GLM and ICA in detecting RSFC patterns has been compared in previous studies, these investigations had some important limitations, including small sample sizes, restricted brain coverage, limited use of advanced noise-reduction methods (e.g., short-channel and physiological signal regression), and most importantly, the exclusive use of HbO signal to determine RSFC and IC selection through visual inspection, which hinders reproducibility.^15,21^ Additionally, the performance of SBA with correlation (SBA-Corr) has not yet been quantitatively compared to other methods.

The present study aims to address these gaps by providing a comprehensive evaluation of SBA and ICA, with a focus on their ability to identify RSFC patterns within the motor network. We investigated both HbO and HbR signals with near whole-head coverage, using standardized preprocessing and artifact-removal methods.

To this end, three variations of SBA were evaluated: SBA using correlation, SBA-GLM, and SBA-GLM with respiratory signal regression (SBA-GLM-Resp). To increase reproducibility and reduce bias, ICA was applied with automated IC selection. Two ICA variations were implemented using different functions to approximate negentropy: (i) ICA that maximizes skewness (ICA-Skew), replicating methodological aspects of Zhang H. et al. (2010) and Behboodi et al. (2019), and (ii) ICA with  ℎ() (ICA-LogCosh), which is described by Hyvärinen et al. (2000) as an optimal and robust approach to approximate negentropy.^28^

Performance was evaluated using ROC curves with functional and anatomical references. The functional reference was defined by the motor task activation map, while the anatomical reference was based on a validated atlas to approximate previous studies.^15,21^ For the anatomical reference, channel-to-region mapping was performed using the fNIRS Optodes’ Location Decider (fOLD) toolbox which relies on validated parcellation methods.^29^ Additionally, each method was evaluated for its ability to generate consistent RSFC patterns in HbO and HbR.

Building on previous findings, we hypothesized that ICA outperforms SBA-GLM in detecting RSFC patterns from HbO signal, as measured by ROC curves,^15,21^ and show higher spatial similarity between HbO and HbR RSFC maps.^15^ Further, we explored which method is more suited for RSFC analysis using fNIRS by systematically comparing SBA and ICA along with their variations. The goal was to provide insights into the strengths and weaknesses of each method and contribute towards their standardization in fNIRS-based RSFC research, ultimately advancing basic neuroscience and promoting clinical translation.

## 2 Methods

### 2.1 Participants & Experimental Tasks

As part of a larger study, 57 healthy subjects were recruited through convenience sampling. Ethical approval was obtained from the Ethics Review Committee of Psychology and Neuroscience at Maastricht University and participants were provided with informed consent and debriefing statements.

Subjects performed a resting state task (5 min) while sitting on a chair, and were instructed to clear their minds, relax, and focus on a fixation cross on a computer screen. This was followed by a bilateral finger-tapping task (5min 50s) with nine motor-action blocks (16s each) separated by jittered fixation periods (15-17s, 1s increments). The finger-tapping run started and ended with a 20s rest period, where a fixation cross was presented on the screen. Participants received 20-Euro compensation upon completion of the study.

### 2.2 Hemodynamic & Physiological Measurements

Hemodynamic signals were recorded using two cascaded NIRSport2 devices and the Aurora v2021.1 software (NIRx Medical Technologies, Berlin). The montage consisted of 134 channels, providing near whole-cortex coverage based on the international 10-10 system, with a sampling rate of 12.6 Hz. It included 32 LED sources, 28 silicon photodiode detectors, and 32 short-distance detectors positioned 8mm from each source (λ1 = 760 nm, λ2 = 850 nm). Optode placement was based on fNIRS fOLD toolbox.^29^ Respiratory signals were simultaneously recorded with the NIRxWings 1.0 system (NIRx Medical Tecnhologies, Berlin).

Fig. 1(a) illustrates the fNIRS cap setup, with red circles representing source optodes, each paired with a short-distance detector shown as a blue ring, and blue circles representing standard-distance detectors. Fig. 1(b) shows the sensitivity profile, computed from Monte Carlo photon propagation simulations. The sensitivity is expressed on a logarithmic scale ranging from 0 to -5, where values near 0 show higher photon sensitivity to tissue changes.

**Fig. 1.**
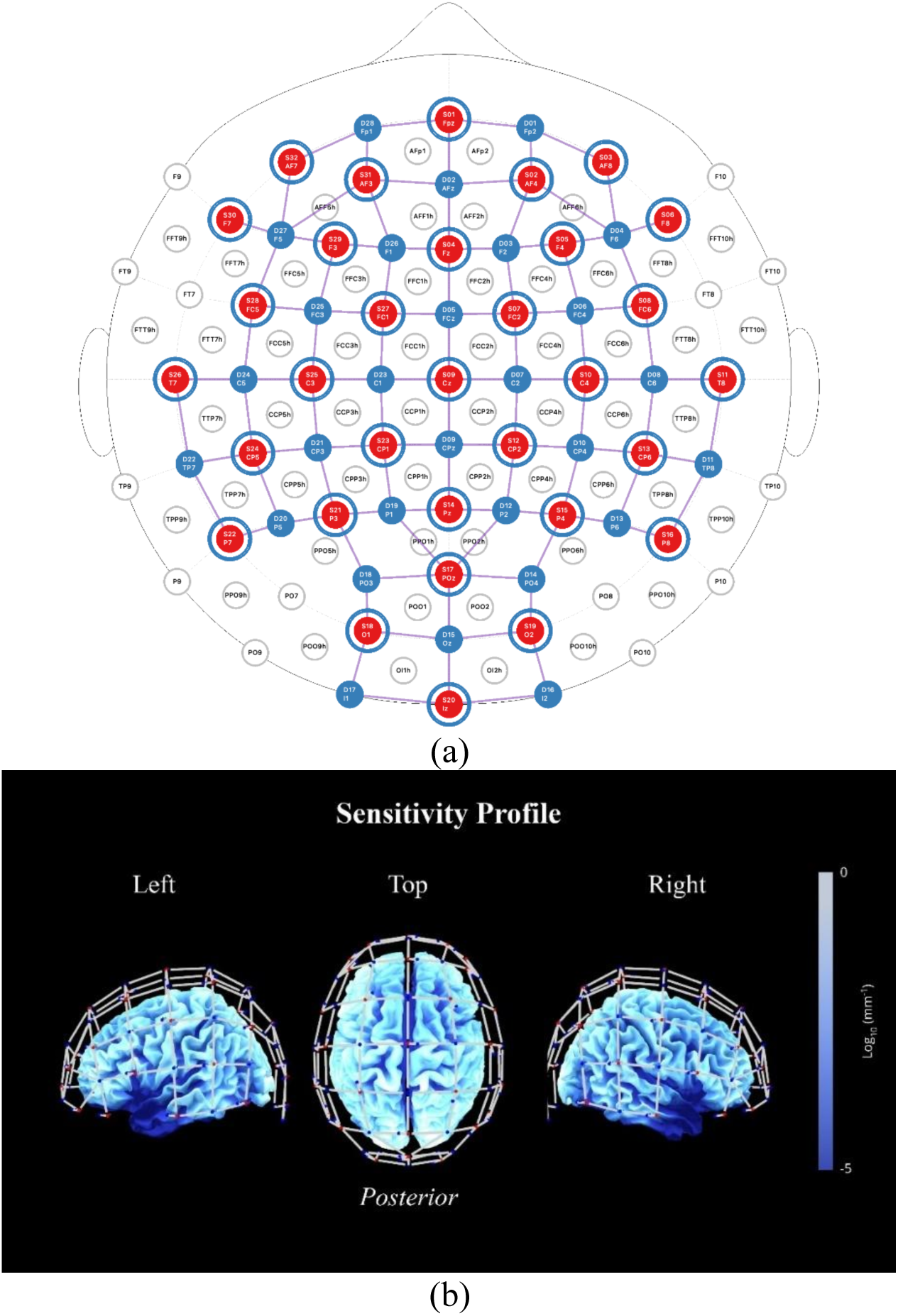
(a) Channel layout showing the spatial arrangement of sources (*N* =32; red circles), standard-distance detectors (*N* = 28; blue circles), and short-distance detectors (*N* = 32; blue rings; 8mm from each source), with optode positions referenced based on the standard EEG landmarks (i.e., Fpz, Fz, etc.) and our custom labeling scheme (e.g., Fpz = S01). (b) Sensitivity profile illustrating photon sensitivity to tissue changes, express on a logarithmic scale ranging from 0 to -5, where values closer to 0 indicate higher sensitivity.

### 2.3 Preprocessing

Data preprocessing and analysis was done using Satori 2.2.4 (Brain Innovation, Maastricht) and Python v3.11.9. For transparency and reproducibility, all code and results are available with each step documented (https://osf.io/whqc9/?view_only=e9fb5c76f21341b29653a161b537d4c9).

#### 2.3.1 Data quality control and exclusion

During data optimization, optode placement was visually inspected and optimized to ensure adequate scalp coupling. Hair was moved away from optodes when necessary, and spring pressure was adjusted to improve signal quality. Signal calibration and online quality monitoring was performed using the Aurora software, and recordings were initiated only after acceptable signal quality was achieved or if 10 minutes of optimizations had passed.

Data exclusion was applied prior to all analyses. Participants were excluded if any of the following criteria were met: (i) insufficient fNIRS data quality, defined as having more than 30% of either standard- or short-distance channels with a scalp coupling index (SCI) below 0.70, where SCI was computed for each channel using Satori after trimming the resting-state time-courses to include only the relevant segments; (ii) incomplete fNIRS runs during either the resting-state or finger-tapping condition; or (iii) poor-quality respiratory recordings during either condition. After applying these criteria, the final sample consisted of 38 participants (*Mage =* 27.71*, SDage =* 10.49; 60.52% female).

#### 2.3.2 Finger-tapping data

Finger-tapping data were preprocessed in the following order: (i) trimming the first 10 seconds to reach steady state;^15–17^ (ii) converting the raw signal to optical density; (iii) rejecting channels with SCI lower than 0.70; (iv) transforming optical density to hemoglobin concentration using the modified Beer-Lambert Law (mBLL); (v) applying a Butterworth bandpass filter of 0.015–0.5 Hz removing contaminating signal from low-frequency noise, respiration, and heart-rate;^15–17,30^ (vi) performing spike removal (iterations = 10; lag = 5s; threshold = 3.5; influence = 0.5; with monotonic interpolation); (vii) applying temporal derivative distribution repair (TDDR) with restoration of high frequencies^30^ (viii) performing GLM-based short-channel regression (using the highest correlated channel); (ix) and *z*-normalization of data.

#### 2.3.3 Resting-state for SBA

For resting-state data in SBA, the same preprocessing steps were applied as in the finger-tapping data, except that the first 20 and last 20 seconds were trimmed to reach steady state,^16,17,20^ a Butterworth band-pass filter of 0.01–0.1 Hz was applied to preserve resting-state signal,^4,11,15,17,31^ and linear trends were removed.^15,20^ Previous fNIRS research using SBA filtered within this range to ensure the signal reflects neuronal activity and is not contaminated by physiological noise (e.g., heart rate, respiration, slow-drift etc.).^14–17,20,21,31^

#### 2.3.4 Resting-state for ICA

As in SBA, for resting state data in ICA the same preprocessing with the finger-tapping data was followed, except for trimming the first 20s and last 20s, detrending the first- and second-order polynomials,^15,21,32^ and applying a broad Butterworth bandpass filter of 0.01–0.2 Hz.^33^ Compared to SBA, the filtering of the resting-state signal is performed by ICA, which is designed to separate different signal sources, including noise. Therefore, a broader input (i.e., 0.01–0.2 Hz) allows for effective identification of resting-state signal and reduces the complexity of selecting the motor relevant component.

### 2.4 Seed Identification

To identify seed channels, a subject-level GLM was performed, with the finger-tapping task as the main predictor,^15–17^ while controlling for *z*-normalized motion parameters (i.e., translation, rotation). The group-level finger-tapping map (hereafter referred to as group motor map) was estimated by performing a one-sample *t-*test on the beta-values of each channel across subjects, with *p*-values being corrected for multiple comparisons using the Benjamini-Hochberg false discovery rate (FDR) method. Channels with mean beta values significantly greater than zero were considered consistently activated across participants during the finger-tapping task. From the resulting group motor map, the channel with the highest absolute *t*-value was selected as the seed for each hemodynamic signal.

### 2.5 RSFC Analysis

#### 2.5.1 SBA-GLM

To estimate RSFC using SBA-GLM, the seed’s time-course was used as a predictor in the GLM for each participant, along with *z-*normalized motion parameters (i.e., translation & rotations) to reduce noise. This produced individual-RSFC maps for each chromophore (HbO and HbR), with beta estimates quantifying the strength of the connectivity between the seed and every other channel. At the group-level, beta values for each channel were compared with a one-sample *t*-test across participants. The null hypothesis stated that the mean beta value for a given channel is equal to zero, indicating no consistent connectivity with the seed. To account for multiple comparisons, the Benjamini-Hochberg FDR correction was applied.

#### 2.5.2 SBA-GLM with respiratory signal

To investigate whether RSFC maps could be improved by accounting for respiratory related fluctuations, a second set of SBA-GLM was performed, in which the same procedure was followed but both the *z*-normalized respiratory signal and seed’s time course were included as predictors in the design matrix (SBA-GLM-Resp), while controlling for *z-*normalized motion parameters (i.e., translation & rotations).

#### 2.5.3 SBA-correlation

Similarly, to estimate RSFC from correlational SBA (SBA-Corr), the seed’s time course was correlated with the time courses of all other channels using Pearson’s *r*, yielding individual-RSFC maps for each chromophore. To compute the group-RSFC, the *r*-values from the individual-RSFC maps were Fisher *z*-transformed and compared using a one sample *t*-test (with Benjamini-Hochberg FDR correction) to identify channels that showed consistent connectivity with the seed across subjects. Fig. 2 summarizes the seed-based analysis pipeline, from preprocessing to individual- and group-level RSFC map estimation.

**Fig. 2.**
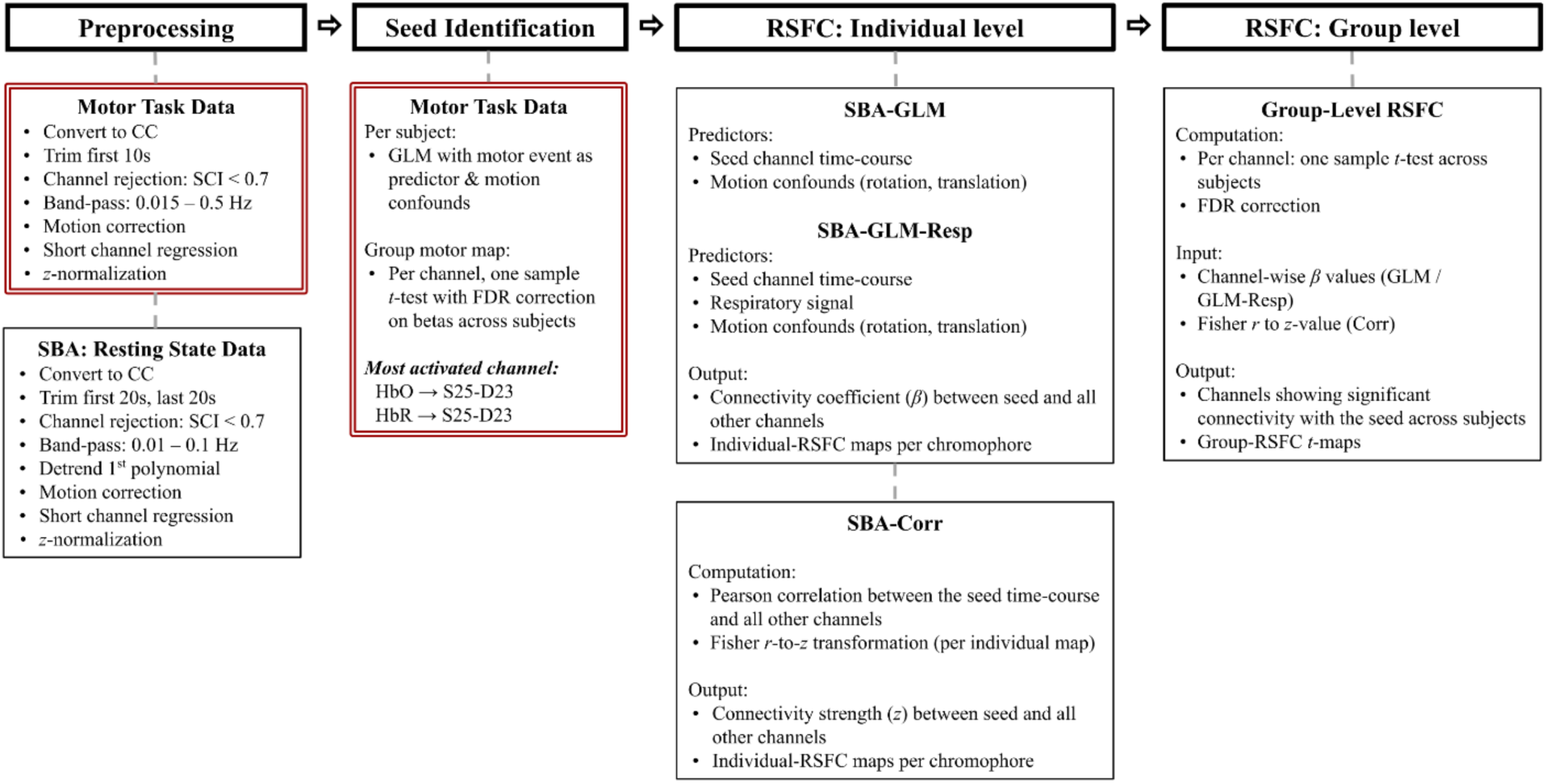
SBA preprocessing and analysis pipeline. The workflow summarizes preprocessing, seed identification, individual-level RSFC estimation, and group-level RSFC estimation for SBA. Finger-tapping data (red outline) and resting-state data (black outline) were preprocessed separately. Seed identification was performed using a subject-level GLM with the motor event as predictor, followed by a one-samples t-test on channel-wise beta values across subjects to generate the group-level motor map. The channel with the highest absolute t-value was selected as the seed. The seed’s time course was used to estimate individual-RSFC maps using three approaches (SBA-GLM, SBA GLM Resp, and SBA-Corr). For SBA-GLM and SBA-GLM-Resp, beta values quantified connectivity between the seed and all other channels. For SBA-Corr, Pearson correlations between the seed and all other channels were computed and Fisher *r*-to-*z* transformed. Group-level RSFC maps for all approaches were obtained using per-channel one-sample *t*-tests with Benjamini-Hochberg FDR correction to identify channels showing significant connectivity with the seed.

#### 2.5.4 ICA

Following and extending the methodology of H. Zhang et al. (2010), a two-stage approach was used to estimate RSFC with ICA.^15^ First, principal component analysis (PCA) was applied to reduce the dimensionality of data, followed by temporal ICA to the PCA-reduced data. A custom Python pipeline (based on scikit-learn 1.7.2) was developed to partially replicate the ICA procedures of H. Zhang et al. (2010), adapting the parameters to match those used in their original MATLAB implementation.

In more detail, PCA was applied separately for HbO and HbR to reduce dimensionality while retaining 99% of the variance, decreasing computational complexity without significant loss of information. Then FastICA was applied on the PCA-reduced data, with the number of components equal to the number of retained principal components. The algorithm was configured to use deflation and a maximum of 10,000 iterations,^15,21^ with tolerance at 10^-6^ to ensure convergence without computational overload. Also, FastICA was performed twice producing two sets of ICA derived RSFC maps, one with the skew function *g*(*u*) = *u*^2^ and one with the *log cosh*(*u*) function, resulting in ICA-Skew and ICA-LogCosh.

To increase validity and reproducibility, the FastICA decomposition was performed multiple times (*N* = 50) per subject, each time with a distinct random seed. Specifically, the random seed was initialized at a fixed value for each run (*seed*= 2025 + *i*, where *i* ∈ {0, 1, …, 49}). This procedure ensures controlled randomness and replicability, avoiding potential bias introduced by relying on a single random seed. Based on prior work, we adopted 50 repetitions as a conservative choice to balance stability with computational feasibility.^34,35^

Therefore, for each participant, 50 sets of ICs were produced per chromophore. To recover the spatial maps of each IC in the original channel space, the PCA component matrix (representing the contribution of each channel to the retained PC) was multiplied with the ICA mixing matrix (representing the contribution of each IC to the PCs). This yielded spatial maps where each IC is expressed as a weighted pattern across channels, which were *z*-score normalized for standardization.

For the selection of motor-related RSFC ICs, a data-driven leave-one-out (LOO) approach was implemented. For each participant and each run, all IC spatial maps were correlated with the group-level motor task activation map computed based on all other participants. Specifically, for each subject, a GLM was performed with the motor task as predictor (and motion confounds) yielding beta spatial maps. To compute the LOO group-level motor map, a one-sample t-test was performed on the beta maps of all other participants, excluding the subject’s own map. This approach ensured that no subject’s motor data contributed to their own IC selection, avoiding circularity and methodological overlap in both IC selection and performance evaluation. Then, the IC with the highest spatial correlation with the LOO motor reference map, was selected, and its sign was aligned to match a reference group motor map (i.e., based on the motor map computed without the first subjects’ data), addressing the sign ambiguity inherent to ICA. Specifically, for the selected IC, if the spatial correlation with the reference group motor map was negative, the sign was flipped. This procedure yielded 50 motor-relevant ICs per participant per chromophore. To enhance stability across runs and reduce noise, the median spatial map was computed across the 50 selected ICs for each participant, resulting in a single IC spatial map for each participant used for group-level analysis.

The individual-RSFC ICA maps were then compared to estimate the group-RSFC map. For each channel and chromophore, a one-sample *t*-test was performed across subjects with the null-hypothesis stating the mean IC weight equals to zero (no consistent activity across subjects), while accounting for the multiple comparison problem with Benjamini-Hochberg FDR correction. In Fig. 3, the ICA analysis pipeline is summarized.

**Fig. 3.**
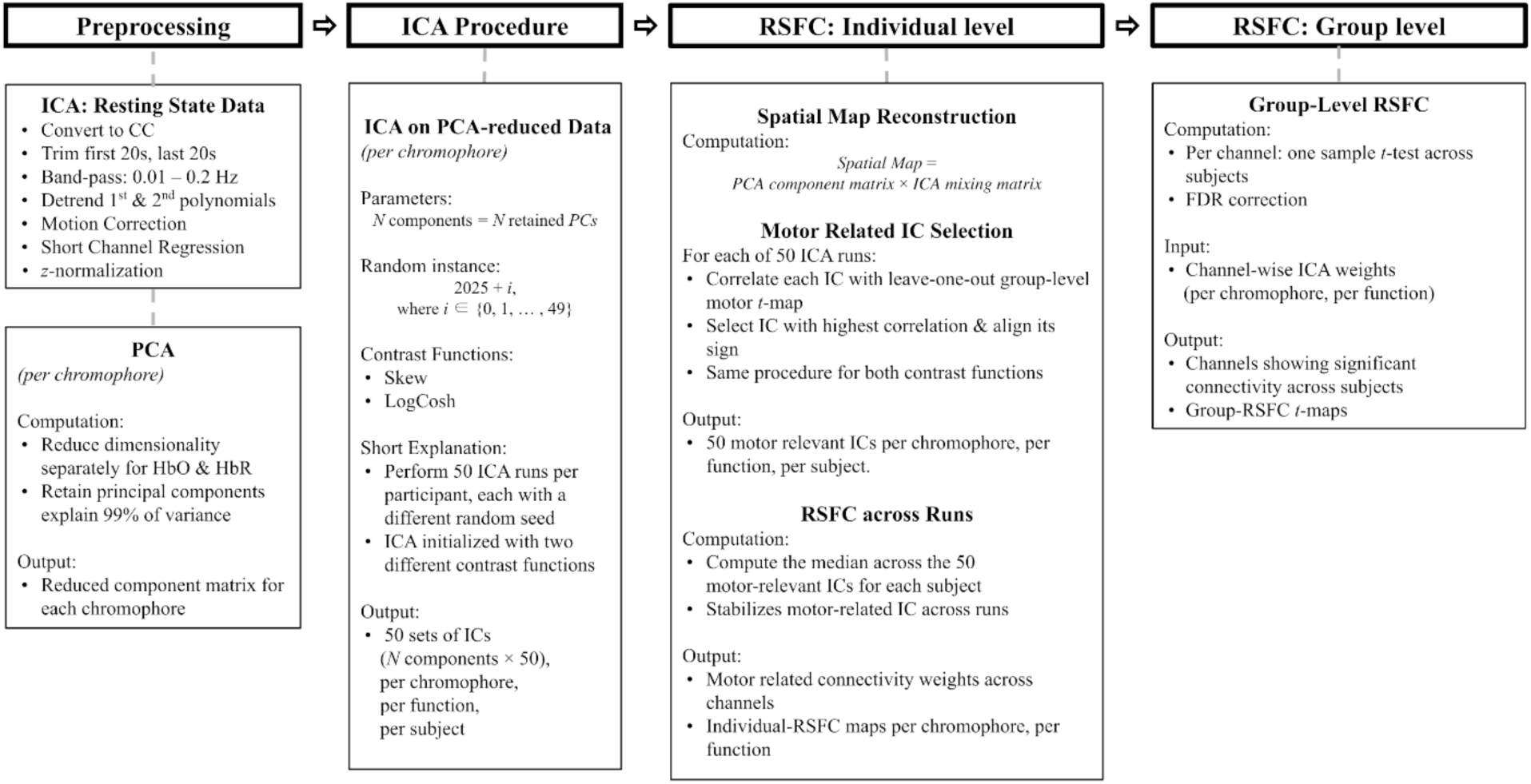
ICA preprocessing and analysis pipeline for RSFC estimation. The workflow summarizes preprocessing, individual-level RSFC estimation, and group-level RSFC estimation for ICA. Resting-state data for ICA were preprocessed similarly to SBA, except for the use of a broader band-pass filter, removal of 1st and 2nd order polynomial trends, and dimensionality reduction using PCA. FastICA was applied to the PCA-reduced data for each chromophore, using two contrast functions (Skew and LogCosh) and 50 repetitions per subject with controlled random seeds (2025 + *i*). Spatial maps of ICs were reconstructed in channel space by combining PCA component matrices with ICA mixing matrices. Per subject, motor-relevant ICs were selected using a data-driven leave-one-out (LOO) approach by correlating each IC spatial map with the corresponding LOO group-level motor map, followed by sign alignment to resolve ICA sign ambiguity. The median spatial map across the 50 selected ICs was computed for each participant to stabilize individual-level RSFC estimates. Group-level RSFC maps were obtained by performing per-channel one-sample t-tests on IC weights across subjects with Benjamini-Hochberg FDR correction, yielding separate group-level RSFC maps for ICA-Skew and ICA-LogCosh.

### 2.6 Performance Evaluation

To assess the ability of each analytical method (i.e., SBA-GLM, SBA-GLM-Resp, SBA-Correlation, ICA-Skew, ICA-LogCosh) in producing reliable RSFC maps, performance was evaluated using (i) ROC curves to quantify how each method predicts motor related activity and (ii) the similarity of HbO and HbR RSFC *t*-maps, based on the assumption that both chromophores reflect activity from the same brain region.

#### 2.6.1 ROC

For the ROC analysis, two binary reference maps were used as golden standards: (i) the group-level motor task activation map derived from the GLM *t-*test procedure (separate map for HbO/HbR; see Sec. 2.4) and (ii) the fOLD atlas, in which channels were mapped to Brodmann areas based on highest specificity, and then the motor relevant channels were grouped together (i.e., B1-B8; see Supplementary Material Sec. S1.1). In the motor activation map, significant channels (*p ≤* 0.05 after FDR correction) were labeled as 1, and non-significant as 0. In the fOLD map, channels corresponding to predefined motor areas were labeled as 1, and all others as 0.

To compute the ROC curves, min-max normalization was applied to the effect size values of each method’s RSFC map.^21^ The performance of each analysis (i.e., SBA-GLM, SBA-GLM-Resp, SBA-Corr, ICA-Skew, and ICA-LogCosh) in predicting motor related activity was then assessed by plotting the true-positive rate against the false-positive rate at varying thresholds.

To evaluate whether the differences between methods were statistically significant, pairwise comparisons of the areas under the curve (AUC) were conducted using DeLong’s test.^36^ This method tests whether the observed differences are due to chance or actual performance difference between methods.

#### 2.6.2 HbO/HbR similarity

To evaluate the similarity of the RSFC patterns in the motor area across chromophores, the spatial correlation of the RSFC *t-*map from HbO and HbR of each analysis was computed using Pearson’s *r*. For the SBA analyses (i.e., SBA-GLM, SBA-GLM-Resp, and SBA-Corr), the *t-*values of the seed channels were masked, due to self-comparison yielding near-infinite values.

## 3 Results

### 3.1 Group-RSFC heatmaps

From the group-RSFC *t-*maps, the number of significant channels after FDR correction between chromophores and across methods are shown in Table S1. Figure 4 shows the group-level RSFC *t*-maps for HbO (left) and HbR (right) across methods.

**Fig. 4.**
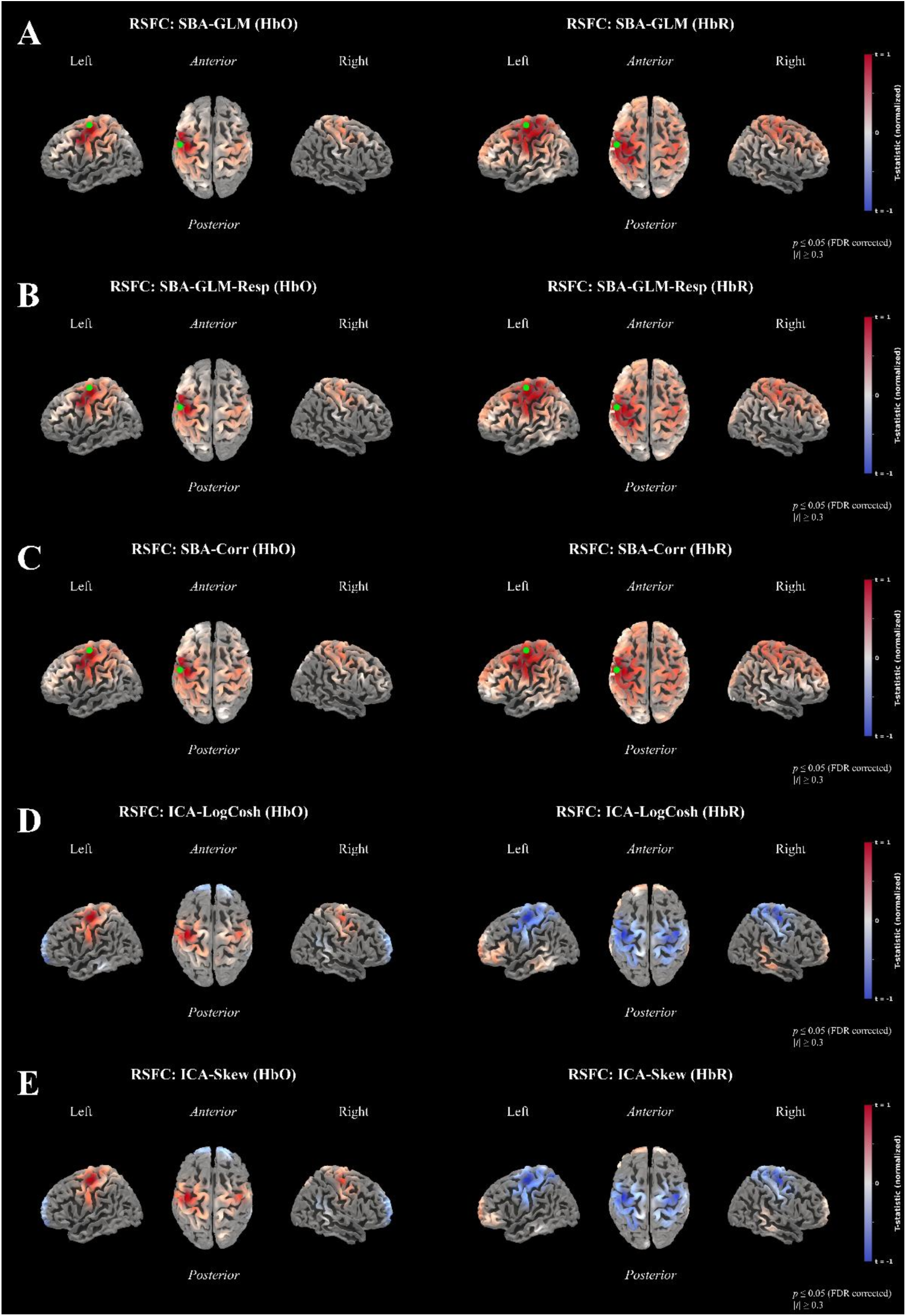
(a) SBA-GLM, (b) SBA-GLM-Resp, (c) SBA-Corr, (d) ICA-LogCosh, and (e) ICA-Skew group-RSFC heatmaps of HbO and HbR. For SBA seed channels are illustrated with a green dot on the left hemisphere, which were capped at the second highest to avoid visualization distortions due to self-comparison. In ICA the negative RSFC motor patterns of HbR are explained by its alignment with the motor task activation map. For visualization purposes, *t*-values were linearly rescaled from -1 to +1, channels with significant group-level connectivity (FDR *p* ≤ 0.05) are shown, and those with |*t*| < 0.3 were masked.

The seed-identification procedure (see Sec. 2.4) revealed a single dominant motor channel that was consistently activated across participants. For both HbO and HbR the channel with the highest absolute *t*-value was S25-D23 in the left hemisphere (HbO: *t* = 12.79, *p* = 3.87 × 10^-13^; HbR: *t* = -15.94, *p* = 3.76 × 10^-16^). This channel was used to estimate the group-RSFC patterns using SBA.

### 3.2 Quantitative Comparison

To assess the performance of each method, ROC analysis was conducted with two reference standards: the fOLD binary map and the finger-tapping activation map, for both HbO and HbR. In ROC analysis, the AUC indicates the similarity between each method’s RSFC results and the reference standard, with higher values reflecting greater similarity.

Using the fOLD binary map as reference, ICA-Skew outperformed the rest for HbO, followed by ICA-LogCosh, SBA-Corr, SBA-GLM-Resp, and SBA-GLM. For HbR, ICA-LogCosh performed best followed by ICA-Skew, SBA-Corr, SBA-GLM and SBA-GLM-Resp. With the motor task activation map binarized at *p* ≤ 0.05 as the reference, for both HbO and HbR, ICA-Skew outperformed the rest, followed by ICA-LogCosh, SBA-Corr, SBA-GLM, and SBA-GLM-Resp (Table 1; Fig 6).

**Table 1.**
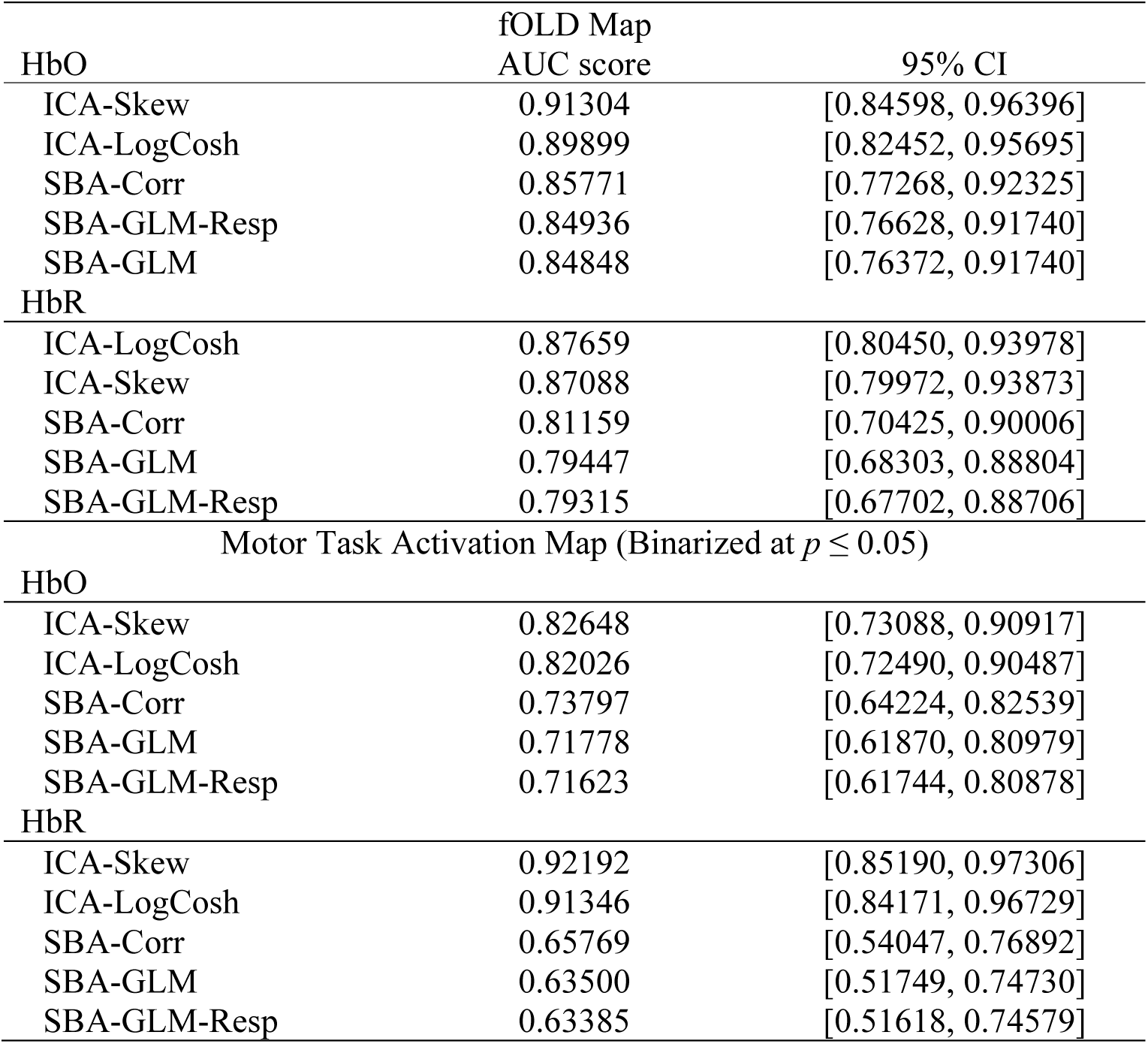
ROC AUC values for RSFC methods, ranked from highest to lowest performance. Results are shown for both HbO and HbR using two reference standards: the fOLD binary motor map and the motor task activation map

| fOLD Map |  |  |
| --- | --- | --- |
| HbO | AUC score | 95% CI |
| ICA-Skew | 0.91304 | [0.84598, 0.96396] |
| ICA-LogCosh | 0.89899 | [0.82452, 0.95695] |
| SBA-Corr | 0.85771 | [0.77268, 0.92325] |
| SBA-GLM-Resp | 0.84936 | [0.76628, 0.91740] |
| SBA-GLM | 0.84848 | [0.76372, 0.91740] |
| HbR |  |  |
| ICA-LogCosh | 0.87659 | [0.80450, 0.93978] |
| ICA-Skew | 0.87088 | [0.79972, 0.93873] |
| SBA-Corr | 0.81159 | [0.70425, 0.90006] |
| SBA-GLM | 0.79447 | [0.68303, 0.88804] |
| SBA-GLM-Resp | 0.79315 | [0.67702, 0.88706] |
| Motor Task Activation Map (Binarized at $p \leq 0.05$) | | |
| HbO |  |  |
| ICA-Skew | 0.82648 | [0.73088, 0.90917] |
| ICA-LogCosh | 0.82026 | [0.72490, 0.90487] |
| SBA-Corr | 0.73797 | [0.64224, 0.82539] |
| SBA-GLM | 0.71778 | [0.61870, 0.80979] |
| SBA-GLM-Resp | 0.71623 | [0.61744, 0.80878] |
| HbR |  |  |
| ICA-Skew | 0.92192 | [0.85190, 0.97306] |
| ICA-LogCosh | 0.91346 | [0.84171, 0.96729] |
| SBA-Corr | 0.65769 | [0.54047, 0.76892] |
| SBA-GLM | 0.63500 | [0.51749, 0.74730] |
| SBA-GLM-Resp | 0.63385 | [0.51618, 0.74579] |

With DeLong’s test, the AUCs for each reference standard across methods and chromophores were tested for differences (see Table 2). In HbR with the fOLD atlas as reference, SBA-Corr significantly outperformed SBA-GLM and SBA-GLM-Resp. With the motor task activation map binarized at *p* ≤ 0.05, for both HbO and HbR signals, SBA-GLM and SBA-GLM-Resp significantly underperformed the rest. In addition, with HbR signal, SBA-Corr significantly underperformed compared to both ICA variations (with a trend towards significance for HbO).

**Table 2.** DeLong’s test comparing AUCs across methods.

| fOLD Map |  |  |  |  |  |
| --- | --- | --- | --- | --- | --- |
| HbO | SBA-GLM | SBA-GLM-Resp | SBA-Corr | ICA-Skew | ICA-LogCosh |
| SBA-GLM | — | $z = 0.4228$,<br>$p = 0.6724$ | $z = 1.1269$,<br>$p = 0.2598$ | $z = 1.8551$,<br>$p = 0.0636$ | $z = 1.3984$,<br>$p = 0.1619$ |
| SBA-GLM-Resp | | — | $z = 1.0178$,<br>$p = 0.3087$ | $z = 1.8528$,<br>$p = 0.0639$ | $z = 1.3942$,<br>$p = 0.1632$ |
| SBA-Corr | | | — | $z = 1.7116$,<br>$p = 0.0869$ | $z = 1.2281$,<br>$p = 0.2194$ |
| ICA-Skew | | | | — | $z = -1.7300$,<br>$p = 0.0836$ |
| ICA-LogCosh |  |  |  |  | — |
| HbR |  |  |  |  |  |
| SBA-GLM | — | $z = -0.7507$,<br>$p = 0.4528$ | $z = \mathbf{1.9721}$,<br>$p = \mathbf{0.0486}$ | $z = 1.4656$,<br>$p = 0.1427$ | $z = 1.6350$,<br>$p = 0.1021$ |
| SBA-GLM-Resp | | — | $z = \mathbf{2.0468}$,<br>$p = \mathbf{0.0407}$ | $z = 1.4816$,<br>$p = 0.1384$ | $z = 1.6513$,<br>$p = 0.099$ |
| SBA-Corr | | | — | $z = 1.1966$,<br>$p = 0.2314$ | $z = 1.3735$,<br>$p = 0.1695$ |
| ICA-Skew | | | | — | $z = 0.7118$,<br>$p = 0.4766$ |
| ICA-LogCosh |  |  |  |  | — |
| Motor Task Activation Map (binarized at $p \leq 0.05$) | | | | | |
| HbO | SBA-GLM | SBA-GLM-Resp | SBA-Corr | ICA-Skew | ICA-LogCosh |
| SBA-GLM | — | $z = -0.7861$,<br>$p = 0.4318$ | $z = \mathbf{2.4516}$,<br>$p = \mathbf{0.0142}$ | $z = \mathbf{2.2616}$,<br>$p = \mathbf{0.0237}$ | $z = \mathbf{2.1590}$,<br>$p = \mathbf{0.0308}$ |
| SBA-GLM-Resp | | — | $z = \mathbf{2.5310}$,<br>$p = \mathbf{0.0113}$ | $z = \mathbf{2.2821}$,<br>$p = \mathbf{0.0224}$ | $z = \mathbf{2.1792}$,<br>$p = \mathbf{0.0293}$ |
| SBA-Corr | | | — | $z = 1.8957$,<br>$p = 0.0580$ | $z = 1.7879$,<br>$p = 0.0737$ |
| ICA-Skew | | | | — | $z = -1.1438$,<br>$p = 0.2527$ |
| ICA-LogCosh |  |  |  |  | — |
| HbR |  |  |  |  |  |
| SBA-GLM | — | $z = -0.6385$,<br>$p = 0.5231$ | $z = \mathbf{2.3510}$,<br>$p = \mathbf{0.0187}$ | $z = \mathbf{5.7254}$,<br>$p < \mathbf{0.0001}$ | $z = \mathbf{5.6404}$,<br>$p < \mathbf{0.0001}$ |
| SBA-GLM-Resp | | — | $z = \mathbf{2.3556}$,<br>$p = \mathbf{0.0185}$ | $z = \mathbf{5.7074}$,<br>$p < \mathbf{0.0001}$ | $z = \mathbf{5.6275}$,<br>$p < \mathbf{0.0001}$ |
| SBA-Corr | | | — | $z = \mathbf{5.4751}$,<br>$p < \mathbf{0.0001}$ | $z = \mathbf{5.3908}$,<br>$p < \mathbf{0.0001}$ |
| ICA-Skew | | | | — | $z = -1.3185$,<br>$p = 0.1873$ |
| ICA-LogCosh |  |  |  |  | — |

Significance at $*p < \mathbf{0.05}$ shown in bold. Positive $z$ indicates higher AUC for the method in the column section.

Additional analyses with stricter binarization thresholds for the motor task activation map were conducted and are reported in Supplementary Materials Sec. S1.3.

### 3.3 HbO/HbR RSFC Patterns Similarity

To assess similarity of RSFC patterns across hemodynamic signals, the linear correlation of the HbO and HbR *t-*maps was computed. Similarity was highest for ICA-Skew (*r* = 0.918, *p* = 3.65 × 10^-42^), followed by ICA-LogCosh (*r* = 0.901, *p* = 4.09 × 10^-38^), SBA-GLM (*r* = 0.8616, *p* = 6.37 × 10^-31^), SBA-GLM-Resp (*r* = 0.8611, *p* = 7.62 × 10^-31^), and SBA-Corr (*r* = 0.84, *p* = 2.07 × 10^-28^).

In Fig. 6, similarity is visualized only for the highest-performing ICA and SBA variations.

## 4 Discussion

In this study, we systematically investigated the ability of SBA and ICA to detect motor-related RSFC patterns with fNIRS data across HbO and HbR signals. The main goal was to advance current methods and work towards their standardization. To evaluate which analysis is more precise, the performance of SBA and ICA was quantitatively assessed with (i) ROC curves and (ii) the similarity of RSFC patterns between HbO/HbR. Overall, ICA is the most suitable analytical approach that identifies consistent RSFC patterns across chromophores. Nonetheless, correlational SBA can be a valid, fast, and low-complexity alternative. Beyond these quantitative metrics, a visual examination of the resulting connectivity maps offers further insights into their ability to detect RSFC networks.

### 4.1 Qualitative Assessment

The heatmaps show that SBA-GLM and SBA-GLM-Resp produce identical RSFC patterns for both chromophores, while SBA-Corr patterns are more diffused compared to the others (Fig. 4). Moreover, across the different SBA approaches, RSFC differs in terms of HbO and HbR. In HbO, there are fewer significant channels, and the motor network is more centralized compared to HbR (Table S1).

For ICA, both LogCosh and Skew reveal two distinct functionally connected networks with opposite activity, one centered in the motor area and another in the frontal region (Fig. 4). In both ICA variations and across chromophores, the motor cortex is clearly identified and localized (Table S1). Nonetheless, there are slight differences in spatial distribution. Regarding the frontal network, across ICA variations there are differences between HbO and HbR (i.e., the identified network in HbR is larger than HbO), indicating that the two signals do not reflect identical spatial patterns of functional connectivity outside the motor region. Overall, ICA produces more consistent RSFC patterns across HbO and HbR. Nonetheless, the motor network is visually identifiable across all methods and chromophores.

### 4.2 Quantitative Assessment

#### 4.2.1 AUCs

These visual findings are supported by ROC analysis, which confirms that RSFC can be accurately detected from both HbO and HbR signals across methods and reference standards (Fig. 5). In previous fNIRS studies, reliable RSFC detection from HbR signal was not possible.^15,21^ With the current experimental setup this limitation was resolved, indicating that functional connectivity can be accurately detected from HbR, which is more relevant to neural activity than HbO.^22^

**Fig. 5.**
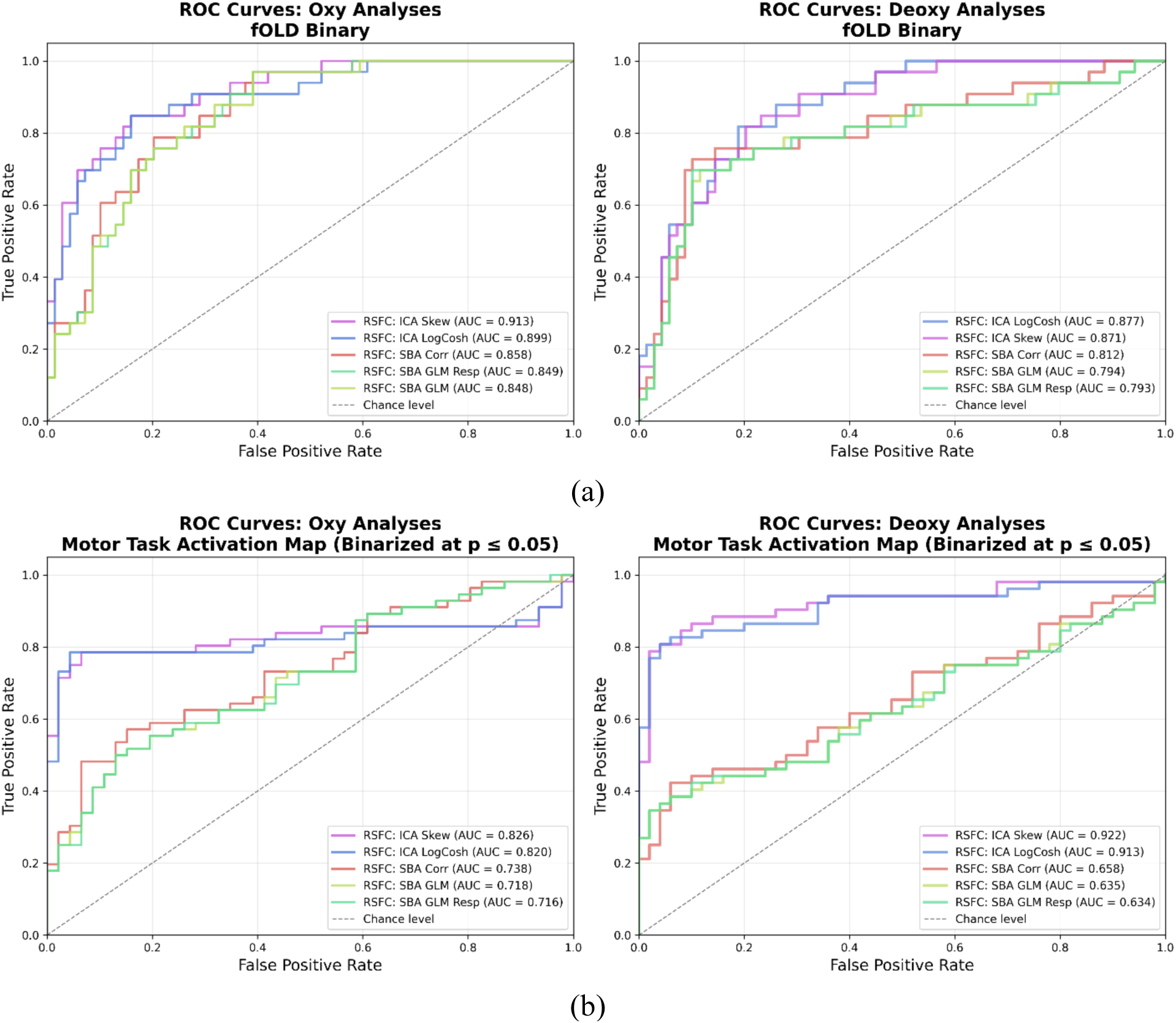
(a) ROC AUC scores of RSFC methods using fOLD and (b) the motor task map (binarized at *p* ≤ 0.05) as reference standards ranked from highest to lowest.

For the ROC analysis, two reference standards were used: an atlas-based map (i.e., a predefined motor map derived from fMRI-to-channel mapping using the fOLD toolbox) and a data-driven functional map (i.e., the activation map from the finger-tapping task). The atlas-based map was chosen to approximate the methodology of H. Zhang et al. (2010) (i.e., localized motor areas using structural MRI from a single subject and mapped the channels to MNI space)^15^ and Behboodi et al. (2019) (i.e., localized motor areas using a 3D-digitizer to map their channels to MNI space using NIR spectroscopy-toolbox)^21^.

The fOLD toolbox provides a standardized approach for channel assignment to anatomical regions. Nonetheless, using structural maps to define functional connectivity has limitations: an anatomical reference represents stable neural structures, while functional connectivity reflects adaptive and changeable neural patterns;^37–39^ and although there is coupling between the two, they are not equivalent^40^ (Supplementary Materials Sec. S1.4, Table S6). Therefore, the motor task activation map was also used as a reference standard, which reflects task-driven functional connectivity rather than structural connectivity.

In terms of AUC values, with either the fOLD map or the motor task activation map (across binarization thresholds) as reference standards, ICA outperformed SBA and its variations across chromophores (Fig. 5, S1, S2). This pattern is in line with our hypothesis for HbO, which predicted higher AUC scores for ICA compared to SBA-GLM, and further demonstrates that the advantage of ICA extends to HbR, across different reference standards, and analysis approaches. These findings are also consistent with Behboodi et al. (2019) and H. Zhang et al. (2010), who used a structural map as the definition of functional connectivity in their ROC analysis.

Additionally, different variations of SBA and ICA were examined. Across chromophores and reference standards, SBA-GLM and SBA-GLM-Resp showed only minor differences in AUC values, indicating limited impact when the respiratory signal is modelled within the GLM framework. In contrast, correlational SBA generally outperformed GLM-based approaches, suggesting that it may be better suited for capturing RSFC patterns. Finally, the two ICA contrast functions showed comparable performance across reference standards, indicating that RSFC estimates were insensitive to the choice of contrast function (Fig. 5, S1, S2).

#### 4.2.2 Statistical Comparison of AUCs

Compared to previous studies,^15,21^ an additional performance assessment was incorporated. DeLong’s test provides a way to statistically compare the AUC values across methods. This approach offers a more robust evaluation of the most effective method for RSFC detection and avoids ambiguous interpretations that arise when AUC CIs overlap.

In more detail, from HbR signal with the fOLD map as reference, correlational SBA showed significantly higher AUCs compared to SBA-GLM and its respiratory variation. This pattern suggests that, under lower signal-to-noise (SNR) conditions typical to HbR signals,^13,22^ GLM-based analyses may show reduced sensitivity in capturing structurally defined functional connectivity (see Table 2). As noted earlier, anatomical maps provide a stable framework of structural connectivity and cannot capture the exact dynamic patterns of functional interactions. As a result, RSFC estimates from all methods fall within the coarse structural boundaries, producing high AUC values across approaches. The general absence of significant differences between methods with DeLong’s test likely reflects this coarse nature of structural maps, which are not sensitive to detect functional distinctions.

Using the functional atlas as a reference, being more relevant to RSFC, revealed widespread performance differences across approaches. Specifically, in both HbO and HbR, ICA-LogCosh, ICA-Skew, and SBA-Corr significantly outperformed SBA-GLM and SBA-GLM-Resp, while SBA-Corr showed a significant disadvantage over ICA variations in HbR and a trend towards significance for HbO (see Table 2). This suggests that although correlational SBA is more effective than GLM-based methods, it is not as powerful as ICA for RSFC detection. Furthermore, under stricter motor map binarization thresholds (i.e., *p* ≤ 0.01 and *p* ≤ 0.001), across chromophores, both ICA variations consistently outperformed all SBA methods, whereas correlational SBA no longer differed from the GLM-based approaches (see Table S3, S5). This pattern illustrates the superiority of ICA in RSFC detection and the relative similarity of SBA methods under a more restrictive definition of RSFC. Interestingly, at the strictest threshold for HbO, SBA-GLM-Resp significantly underperformed SBA-GLM, suggesting that regressing out the respiratory signal may remove components relevant to RSFC when the positive class is narrowly defined. However, this effect was not consistent across all thresholds or reference maps, indicating that its impact cannot be generalized.

Overall, DeLong’s test provided statistical evidence that ICA-based approaches consistently outperformed SBA methods across chromophores and threshold levels, particularly when evaluated against a functional atlas. While correlational SBA surpassed the GLM-based approaches under moderate thresholds, ICA remained the most reliable method for RSFC detection, with clear advantage in lower SNR conditions. These results highlight the ability of ICA in extracting functional connectivity accurately, and the benefit of using statistical comparisons, such as DeLong’s test, to validate differences in model performance. The findings further suggest that ICA is more effective in capturing functional connectivity patterns that remain stable even when the definition of connectivity becomes more selective. An alternative explanation is that stricter thresholding reduced the size of the positive reference set, which may differentially penalize SBA, as it tends to produce more diffused spatial patterns.

#### 4.2.3 Spatial similarity of HbO/HbR RSFC maps

While ROC analysis measures each method’s ability to detect RSFC patterns, the spatial similarity between HbO and HbR RSFC maps evaluates the consistency of RSFC patterns across chromophores for each method. Both HbO and HbR signals reflect the same underlying neural activity, which should be captured by the method used to estimate RSFC. Across methods, the spatial similarity between chromophores was generally high, but SBA approaches showed lower overlap than ICA methods (Fig. 6), consistent with previous findings^15^ and in line with the second part of our hypothesis, which predicted that ICA would show higher similarity in RSFC patterns between HbO and HbR.

**Fig. 6.**
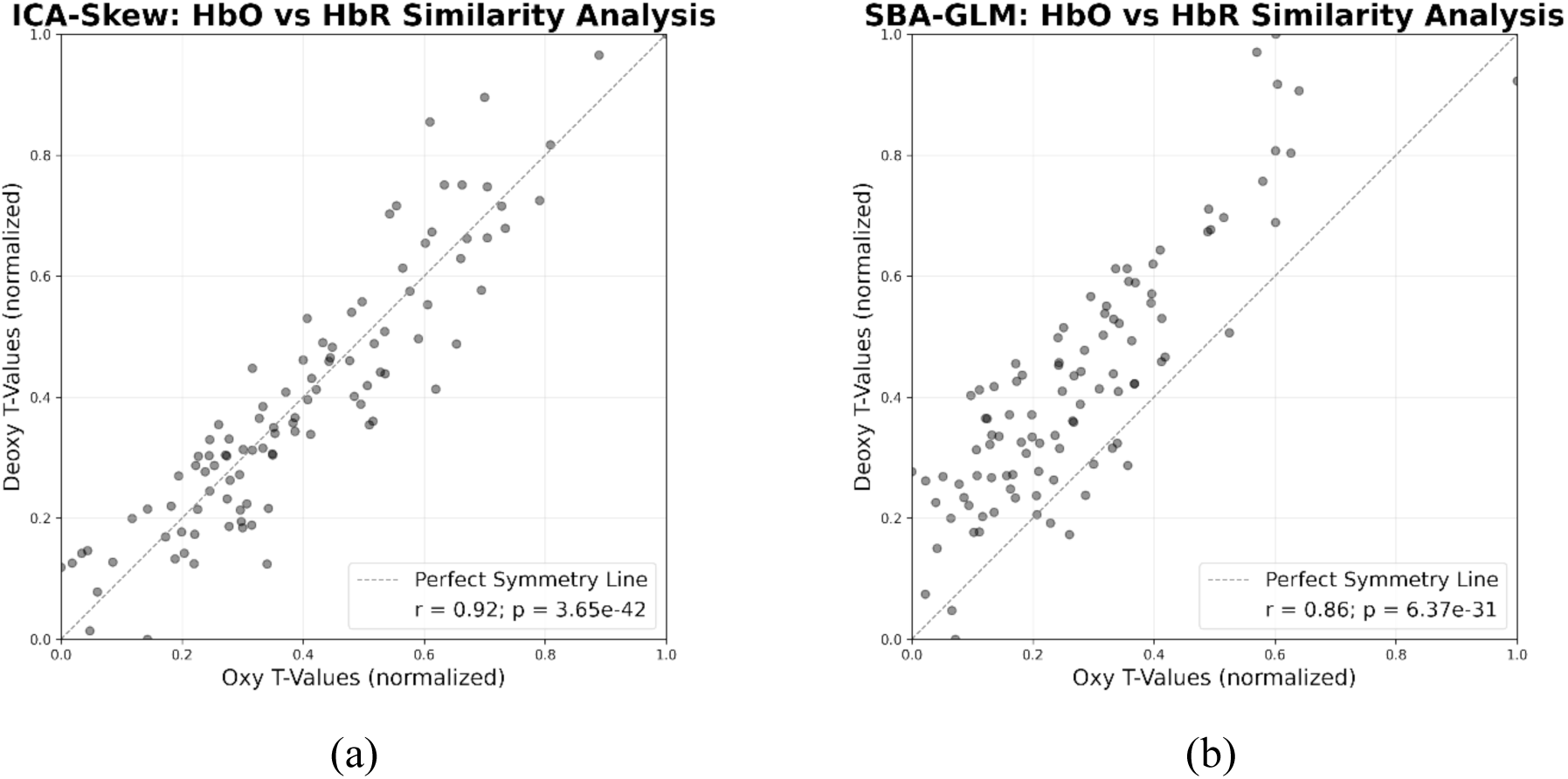
Similarity of motor RSFC patterns between HbO/HbR for (a) ICA-Skew and (b) SBA-GLM. For visualization purposes only, the *t*-values were normalized in the range of 0 to 1

Specifically, ICA-Skew had the highest similarity, closely followed by ICA-LogCosh; indicating that both contrast functions extract similar signal patterns from HbO and HbR. However, it should be noted that there were certain methodological differences compared to previous research. ICA was performed on the whole-head, which identified an additional frontal network. This network showed differences in overlap between HbO and HbR (Fig. 4). Although this may have slightly reduced the spatial similarity of the HbO/HbR RSFC patterns in ICA, it did not drive the effect, illustrating the robustness of ICA in extracting the relevant motor network from both chromophores effectively.

Overall, the performance evaluation in the present study demonstrates that both HbO and HbR signals are informative for RSFC mapping and can yield comparable and accurate patterns of functional connectivity. Previous studies have primarily focused on HbO signals when evaluating the ability of SBA and ICA to detect RSFC patterns with ROC analysis.^15,21^ Extending this prior work, our results show that among the tested approaches, ICA appears to provide the most stable, robust, and reliable estimates of functional connectivity across both chromophores. Nonetheless, this does not diminish the utility of SBA approaches, as correlational SBA proved to be an efficient and accurate alternative for RSFC detection. These findings suggest that HbR contains meaningful functional connectivity information comparable to HbO and that the choice of analytical approach plays a critical role in determining RSFC reliably for both chromophores.

### 4.3 Strengths & Limitations

A central aspect of the current study was the use of both HbO and HbR to detect connectivity patterns. With the present methodological considerations, RSFC could be reliably identified from HbR with both ICA and SBA. Unlike previous studies, which had restricted head-coverage and relied on manual and subjective selection of ICs, the current approach involved a near whole-head setup combined with additional preprocessing, providing sufficient spatial information for ICA to extract distinct functional networks.^28^

Importantly, ICA algorithms are inherently stochastic, such that different random initializations may converge to different local minima and yield different results, even when applied on the same data.^34,35^ To address this, ICA was repeated 50 times with different initial seeds and results were combined.^34^ Previous work on fMRI suggests that stability for FastICA is achieved with 20 to 60 repetitions, depending on the dataset,^35^ while comparable stability for other non-deterministic models (i.e., Hidden Markov Model) has been reported with approximately 50 repetitions.^34^ Moreover, IC selection was fully automated using a reproducible, data-driven leave-one-out approach, reducing ambiguity in the identification of motor-relevant components. Collectively, repeating ICA with multiple initializations, selecting motor-relevant ICs with a data-driven approach, and combining the results across runs, provided a robust and reliable estimate of the motor RSFC network. These methodological considerations were not evident in prior work.

Furthermore, this study is the first to use a functional map as a reference standard to evaluate SBA and ICA, which is more relevant to RSFC than a structural map. The use of two different RSFC definitions yielded somewhat different results, highlighting the importance of reference selection on RSFC outcomes. Furthermore, the comparison of SBA-Correlation with SBA-GLM and ICA has not been performed previously and revealed important findings for future research in RSFC. SBA-Correlation proved to be an effective alternative to SBA-GLM using fNIRS. Additionally, using DeLong’s test to compare ROC curves, which was also not implemented in earlier work, strengthened the interpretability of performance differences across methods. Lastly, using current preprocessing standards, this study updates our understanding of the relative strengths and limitations of each approach for RSFC analysis.

Moving on, several limitations should be acknowledged. First, the analysis focused on the motor cortex, and it remains unclear whether ICA outperforms SBA in other networks or whether structural atlases versus data-driven functional references produce different results outside the motor system. Secondly, in ICA a single IC was selected per participant, but the combination of multiple ICs may better represent the network of interest and improve overall performance. Thirdly, it is unclear whether ICA’s effectiveness was driven by the near whole-head set up, which provided more information to detect the functional network of interest. Lastly, in SBA seed identification was performed on the group-level, but performance could potentially be improved by defining seeds individually (see Supplementary Materials Sec. S1.5).

### 4.4 Future Considerations

Addressing the above limitations could refine our understanding of RSFC with fNIRS. Future studies should consider larger sample sizes, examine the performance of ICA and SBA across multiple brain networks, with different reference standards to assess their generalizability. Moreover, it is essential to ensure that reference standards accurately represent functional connectivity to allow valid method comparisons. This could be achieved through simultaneous fMRI-fNIRS recordings or by using standardized functional brain maps aligned with the fNIRS channel layout. Methodological improvements may also increase the accuracy and consistency of RSFC mapping. For ICA this could involve (i) combining multiple ICs to better represent the network of interest and (ii) using lower convergence thresholds with different contrast functions. Further, SBA could be improved by implementing individualized seed selection based on functional localizers.

From our current findings, both SBA and ICA can be adapted for practical applications in group-RSFC estimation. Correlational SBA could prove beneficial for quick and accurate assessment of functional connectivity. Moreover, in settings where a localizer task cannot be performed, both SBA and ICA can still be used to estimate functional connectivity with a potential tradeoff in accuracy. For instance, in SBA seed selection could be performed manually through a structural atlas such as fOLD, while ICA’s automatic IC selection can be guided by functional atlases derived from fMRI literature and adapted to the fNIRS space.

Although this was not within the scope of the present study, visual inspection of the heatmaps in Fig. S4 indicates that the motor network could be identified at the single-subject level with both ICA and SBA. The ability to reliably detect networks at the subject-level is critical for the translation of fNIRS-based RSFC analyses to clinical practice. At present, repeated assessment of functional connectivity using fMRI is often impractical due to high costs,^6,41^ limited scanner availability,^42^ and additional logistical constraints.^23,41,42^ In this context, fNIRS offers a promising alternative for longitudinal RSFC monitoring, with practical advantages over fMRI and a clear sensitivity in detecting group-RSFC patterns. Future work should focus on evaluating the reliability of fNIRS for capturing individualized RSFC estimates. Such efforts would further support the clinical translation of fNIRS, ideally through validation studies employing simultaneous fNIRS-fMRI measurements.

Beyond ICA and SBA, the field of RSFC with fNIRS could benefit from exploring and comparing alternative analytical methods. Graph theory has been increasingly used to estimate RSFC^3,4,20^ but has not yet been systematically compared to SBA and ICA, possibly due to methodological differences. Furthermore, machine-learning techniques have shown great potential in neuroscience but remain largely unutilized in fNIRS-based RSFC.^43^ ANN and CNN represent promising alternatives for RSFC estimation, but further research is needed to standardize their application.^21^

## 5 Conclusion

This study provides a comprehensive evaluation of SBA and ICA for RSFC detection based on fNIRS recordings. The findings demonstrate that ICA is generally more effective in identifying functional connectivity across chromophores, and should be preferred for RSFC estimation, particularly when connectivity patterns from both chromophores are required. In contrast, SBA requires a seed region, typically obtained through a localizer task. When such information is available, ICA is well positioned to estimate RSFC patterns of the region of interest accurately, as the relevant component can be selected through an unambiguous, data-driven approach. Even in the absence of localizer data, previous studies have shown that visual inspection of ICs can reliably identify the relevant component, providing better RSFC estimation than SBA with visually selected seeds. However, performing ICA correctly can be complex, which may pose a challenge for some users. In such cases, correlation SBA remains a viable and efficient alternative. Overall, this work refines current analytical approaches and supports efforts to standardize fNIRS-based RSFC methodology, which is a critical step toward advancing both basic neuroscientific research and clinical application.

## 7 Code, Data, and Materials Availability

For transparency and replicability, all code used in this study along with the generated results and analysis outputs are available in Open Science Framework (https://osf.io/whqc9/?view_only=e9fb5c76f21341b29653a161b537d4c9). A README file is also included to facilitate navigation through the folders and clarify the structure and usage of the provided materials.

The original fNIRS files (.snirf) cannot be shared publicly at this stage because they are part of an unpublished dataset currently under review for publication. This data can be made available upon request and with permission from the corresponding author.

## Acknowledgments

This work was supported by The Netherlands Organization for Scientific Research (NWO; Vidi-Grant No. VI.Vidi.191.210 to BS).

During the preparation of this work, ChatGPT was used to edit English phrasing and grammar in the text. Following the use of this tool, we reviewed and edited the content as necessary and take full responsibility for the content of this publication.

## Author Biographies

Biographies and photographs for the authors are not available.

---

**Fig. S1** ROC AUC scores of RSFC methods using the motor task map binarized at *p* ≤ 0.01 ranked from highest to lowest.

**Fig. S2** ROC analysis of RSFC methods using the motor task map binarized at *p* ≤ 0.001 ranked from highest to lowest.

**Fig. S3** The top row shows histograms of the 10 most frequently selected channels for HbO (left) and HbR (right) across subjects (*N* = 38). The full montage (bottom right) provides a reference for the complete channel layout, illustrating the spatial arrangement of all channels across the scalp. In the cropped montage (middle), the two most representative motor seeds for each signal are highlighted with orthogons: red orthogons indicate HbO seeds, and blue orthogons indicate HbR seeds.

**Fig. S4** (a) SBA-Corr and (b) ICA-LogCosh RSFC heatmaps of HbO and HbR from a single subject at random. For SBA, the seed channel (S25-D23) is illustrated with a green dot on the left hemisphere. For visualization purposes, the *r*-value of the seed channel was capped at the second highest (to avoid visualization distortions), and then all *r*-values were divided by the absolute maximum r to produce a normalized map ranging from -1 to +1, comparable to ICA. For ICA, the subject’s RSFC map was obtained by taking the median across 50 ICA runs, and then the channel weights were linearly rescaled from -1 to +1. For both SBA and ICA, the values below 0.3 were masked.

**Table S1** Number of significant channels in the group-RSFC maps across methods and chromophores.

**Table S2** ROC AUC values for RSFC methods, ranked from highest to lowest performance. Results are shown for both HbO and HbR using the motor task activation map binarized at *p* ≤ 0.01.

**Table S3** DeLong’s test comparing AUCs across methods with the motor map binarized at *p* ≤ 0.01.

**Table S4** ROC AUC values for RSFC methods, ranked from highest to lowest performance. Results are shown for both HbO and HbR using the motor task activation map binarized at *p* ≤ 0.001.

**Table S5** DeLong’s test comparing AUCs across methods with the motor map binarized at *p* ≤ 0.001.

**Table S6** Spatial correlation between the fOLD binary map (structural) and the binarized motor-task activation maps (functional) at different binarization thresholds.

