## Supplementary Materials for "Independent Component Analysis Outperforms Seed-Based Approach in Detecting fNIRS-based Resting-State Functional Connectivity"

### Supplementary Material

#### *S1.1 fOLD Binarization Process*

The channel-to-region mapping was determined using the fOLD toolbox (https://github.com/nirx/fOLD-public). The program was configured to have forced symmetry (i.e., equal specificity for channels in mirrored locations), and the channels were mapped to Brodmann areas using the 10-10 system. Then the extracted labels were converted to match our study's labeling system (e.g., from Fpz-Fp2 to S1-D1). Finally, Brodmann areas were grouped into broader categories based on the BrainTutor program (Brain Innovation, Maastricht). Broadman areas corresponding to the primary motor (i.e., B4), somatosensory (i.e., B1, B2, and B3), secondary somatosensory (i.e., B5 and B7), and premotor cortex (B6 and B8) were merged to represent the motor-related channels used in the atlas-based reference standard in the present study. Please refer to the OSF folder (Raw_Results/91_MULPA fOLD areas) containing the raw output from the fOLD toolbox, with each processing step documented and illustrated.

#### *S1.2 Number of Significant Channels in Group-RSFC Maps*

**Table S1** Number of significant channels in the group-RSFC maps across methods and chromophores.

| Method | Chromophores | |
|---|---|---|
| | HbO | HbR |
| SBA-GLM | 66 | 75 |
| SBA-GLM-Resp | 65 | 76 |
| SBA-Corr | 61 | 76 |
| ICA-LogCosh | 66 | 69 |
| ICA-Skew | 65 | 64 |

*Note.* Significance at $p < 0.05$ (FDR corrected). Total number of channels (excluding short-distance detectors): 102

#### *S1.3 ROC Analysis with Stricter Thresholding*

An additional analysis investigated how stricter thresholding would affect the accuracy of each method to predict the motor task activation map.

##### *S1.3.1 Binarization at $p \leq 0.01$*

With binarization of the motor map at $p \leq 0.01$, in HbO, ICA-LogCosh outperformed the rest, followed by ICA-Skew, SBA-Corr, SBA-GLM, and SBA-GLM-Resp. In HbR, ICA-LogCosh and ICA-Skew yielded identical AUC values and outperformed all other methods, followed by SBA-Corr, SBA-GLM-Resp, and SBA-GLM (SBA-GLM-Resp and SBA-GLM also exhibited identical AUC values) (Table S2, Fig. S1)

DeLong's test showed that for both HbO and HbR, ICA-LogCosh and ICA-Skew significantly outperformed all SBA methods (see Table S3).

**Table S2** ROC AUC values for RSFC methods, ranked from highest to lowest performance. Results are shown for both HbO and HbR using the motor task activation map binarized at $p \leq 0.01$.

| Motor Task Activation Map (Binarized at $p \leq 0.01$) | | |
|---|---|---|
| HbO | AUC score | 95% CI |
| ICA-LogCosh | 0.89597 | [0.80429, 0.96577] |
| ICA-Skew | 0.89556 | [0.80648, 0.96545] |
| SBA-Corr | 0.75329 | [0.65012, 0.85381] |
| SBA-GLM | 0.75082 | [0.64748, 0.85123] |
| SBA-GLM-Resp | 0.74671 | [0.64186, 0.84788] |
| HbR | | |
| ICA-LogCosh | 0.96230 | [0.92749, 0.98769] |
| ICA-Skew | 0.96230 | [0.92780, 0.98800] |
| SBA-Corr | 0.75357 | [0.64305, 0.84649] |
| SBA-GLM-Resp | 0.74206 | [0.62711, 0.84097] |
| SBA-GLM | 0.74206 | [0.63020, 0.84094] |

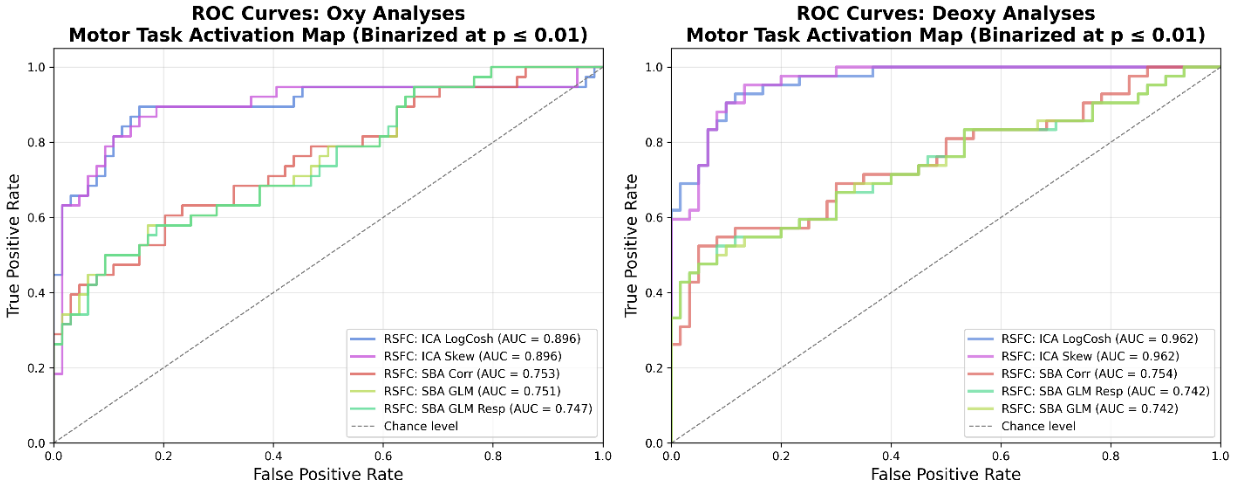


**Fig. S1** ROC AUC scores of RSFC methods using the motor task map binarized at $p \leq 0.01$ ranked from highest to lowest.

**Table S3** DeLong's test comparing AUCs across methods with the motor map binarized at $p \leq 0.01$.

| Motor Task Activation Map (Binarized at $p \leq 0.01$) | | | | | |
|---|---|---|---|---|---|
| HbO | SBA-GLM | SBA-GLM-Resp | SBA-Corr | ICA-Skew | ICA-LogCosh |
| SBA-GLM | – | $z$ = -1.8766, $p$ = 0.0606 | $z$ = 0.2657, $p$ = 0.7905 | **$z$ = 3.4597, $p$ = 0.0005** | **$z$ = 3.5138, $p$ = 0.0004** |
| SBA-GLM-Resp | | – | $z$ = 0.6573, $p$ = 0.5110 | **$z$ = 3.5298, $p$ = 0.0004** | **$z$ = 3.5759, $p$ = 0.0003** |
| SBA-Corr | | | – | **$z$ = 3.4950, $p$ = 0.0005** | **$z$ = 3.5597, $p$ = 0.0004** |
| ICA-Skew | | | | – | $z$ = 0.0636, $p$ = 0.9493 |
| ICA-LogCosh | | | | | – |
| HbR | | | | | |
| SBA-GLM | – | $z$ = 0, $p$ = 1 | $z$ = 1.2686, $p$ = 0.2046 | **$z$ = 4.3989, $p$ < 0.0001** | **$z$ = 4.4083, $p$ < 0.0001** |
| SBA-GLM-Resp | | – | $z$ = 1.2307, $p$ = 0.2184 | **$z$ = 4.3865, $p$ < 0.0001** | **$z$ = 4.3969, $p$ < 0.0001** |
| SBA-Corr | | | – | **$z$ = 4.3320, $p$ < 0.0001** | **$z$ = 4.3438, $p$ < 0.0001** |
| ICA-Skew | | | | – | $z$ = 0, $p$ = 1 |
| ICA-LogCosh | | | | | – |

Significance at **$*p < 0.05$** shown in bold. Positive $z$ indicates higher AUC for the method in the column section.

*S1.3.2 Binarization at $p \leq 0.001$*

When the motor task map was binarized at $p \leq 0.001$, for both chromophores, ICA-Skew outperformed the rest, followed by ICA-LogCosh, SBA-Corr, SBA-GLM, and SBA-GLM-Resp (SBA-Corr and SBA-GLM yielded identical AUC scores for HbR) (Table S4, Fig. S2).

DeLong's test showed that for both HbO and HbR, ICA-LogCosh and ICA-Skew significantly outperformed the rest, and for HbO particularly, SBA-GLM significantly outperformed SBA-GLM-Resp (Table S5).

**Table S4** ROC AUC values for RSFC methods, ranked from highest to lowest performance. Results are shown for both HbO and HbR using the motor task activation map binarized at $p \leq 0.001$.

| Motor Task Activation Map (Binarized at $p \leq 0.001$) | | |
|---|---|---|
| HbO | AUC score | 95% CI |
| ICA-Skew | 0.92631 | [0.84120, 0.98178] |
| ICA-LogCosh | 0.92395 | [0.84002, 0.98083] |
| SBA-Corr | 0.81153 | [0.70849, 0.90136] |
| SBA-GLM | 0.80822 | [0.70510, 0.89754] |
| SBA-GLM-Resp | 0.80350 | [0.69810, 0.89390] |
| HbR | | |
| ICA-Skew | 0.96268 | [0.92349, 0.98844] |
| ICA-LogCosh | 0.95276 | [0.90848, 0.98353] |
| SBA-GLM | 0.78649 | [0.67747, 0.88704] |
| SBA-Corr | 0.78649 | [0.68116, 0.88409] |
| SBA-GLM-Resp | 0.78602 | [0.67736, 0.88662] |

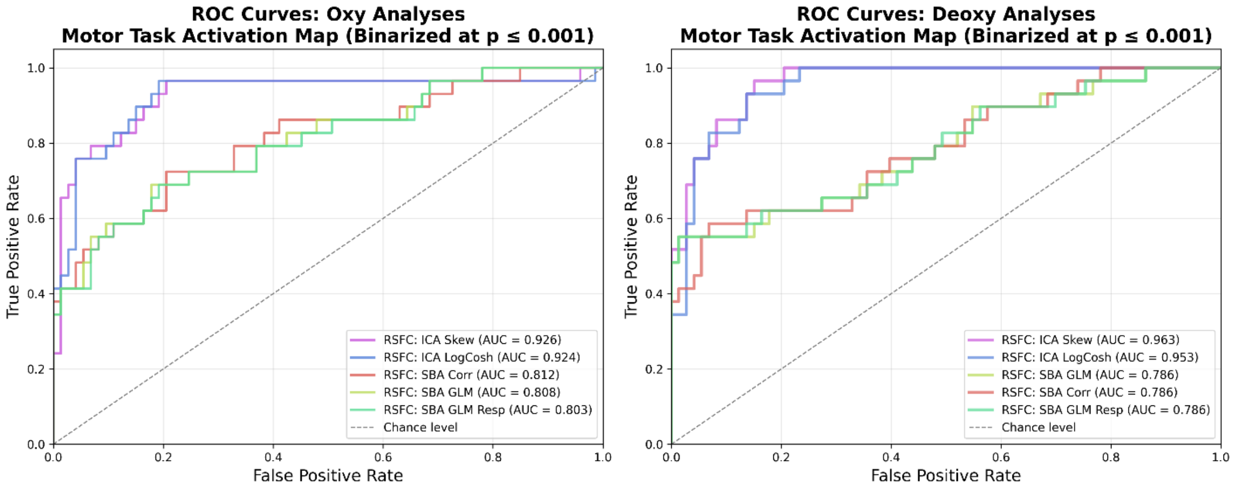


**Fig. S2** ROC analysis of RSFC methods using the motor task map binarized at $p \leq 0.001$ ranked from highest to lowest.

**Table S5** DeLong's test comparing AUCs across methods with the motor map binarized at $p \leq 0.001$.

| Motor Task Activation Map (binarized at $p \leq 0.001$) | | | | | |
|---|---|---|---|---|---|
| HbO | SBA-GLM | SBA-GLM-Resp | SBA-Corr | ICA-Skew | ICA-LogCosh |
| SBA-GLM | – | **$z$ = -2.1464, $p$ = 0.0318** | $z$ = 0.3514, $p$ = 0.7253 | **$z$ = 2.8552, $p$ = 0.0043** | **$z$ = 2.7983, $p$ = 0.0051** |
| SBA-GLM-Resp | | – | $z$ = 0.7915, $p$ = 0.4286 | **$z$ = 2.9321, $p$ = 0.0033** | **$z$ = 2.8736, $p$ = 0.0040** |
| SBA-Corr | | | – | **$z$ = 2.7539, $p$ = 0.0059** | **$z$ = 2.7064, $p$ = 0.0068** |
| ICA-Skew | | | | – | $z$ = -0.2822, $p$ = 0.7778 |
| ICA-LogCosh | | | | | – |
| HbR | | | | | |
| SBA-GLM | – | $z$ = -0.1866, $p$ = 0.8520 | $z$ = 0, $p$ = 1 | **$z$ = 3.5826, $p$ = 0.0003** | **$z$ = 3.3275, $p$ = 0.0009** |
| SBA-GLM-Resp | | – | $z$ = 0.0467, $p$ = 0.9627 | **$z$ = 3.5779, $p$ = 0.0003** | **$z$ = 3.3232, $p$ = 0.0009** |
| SBA-Corr | | | – | **$z$ = 3.7145, $p$ = 0.0002** | **$z$ = 3.4605, $p$ = 0.0005** |
| ICA-Skew | | | | – | $z$ = -1.6497, $p$ = 0.0990 |
| ICA-LogCosh | | | | | – |

Significance at ***$p$ < 0.05** shown in bold. Positive $z$ indicates higher AUC for the method in the column section.

#### *S1.4 Structural Maps Versus Functional Maps to Define RSFC Networks*

The fOLD binary map, being an anatomical reference, does not necessarily reflect functional connectivity. Therefore, the motor-task activation map was also used as a more relevant definition of RSFC. Although structure and function are related, they are not equivalent, and in Table S6 this is illustrated. The structural map (fOLD) shows spatial correspondence with the functional map (motor-map), but no complete overlap. Therefore, anatomical maps should be used with caution as ground truth in RSFC analyses.

**Table S6** Spatial correlation between the fOLD binary map (structural) and the binarized motor-task activation maps (functional) at different binarization thresholds.

| Binarization threshold | HbO | | HbR | |
|---|---|---|---|---|
| | Pearson's *r* | *p* | Pearson's *r* | *p* |
| 0.05 | 0.416 | **$1.35 \times 10^{-5}$** | 0.468 | **$6.82 \times 10^{-7}$** |
| 0.01 | 0.421 | **$1.06 \times 10^{-5}$** | 0.486 | **$2.25 \times 10^{-7}$** |
| 0.001 | 0.447 | **$2.51 \times 10^{-6}$** | 0.354 | **$2 \times 10^{-4}$** |

*Note.* Significance at ***p* < 0.05** in bold

#### *S1.5 Seed Selection on Individual Basis*

Our primary aim was to compare analytical methods while following the methodology of previous studies. To this end, seed identification was performed as described in previous literature,[15,16,20,44] with the seed channel defined based on group-level results. Specifically, after obtaining the group motor map from the bilateral finger-tapping task, the most activated channel was defined as the seed. This seed was then used in SBA to derive individual-RSFC maps, which were compared using a *t*-test to obtain the group-RSFC maps.

An alternative approach would be to identify the seed individually for each participant by performing an individual-level GLM of the motor task and selecting the most activated channel as the seed. This participant-specific seed could be used in SBA to estimate individual-RSFC networks and then the group-RSFC map. This method accounts for inter-individual variability in cortical organization and cap placement.

As shown in Figure S3, there is variability in the location of the motor seed on individual basis. In HbO, 17 unique channels were identified as seeds with the most common one being S10-D7 in 8 participants. Similarly, in HbR, 20 unique channels were defined as seeds with S25-D23 appearing as optimal for 9 subjects. This variation indicates that a fixed seed defined at the group-level does not necessarily provide the most optimal RSFC estimates per individual, and in effect it may reduce the performance of SBA.

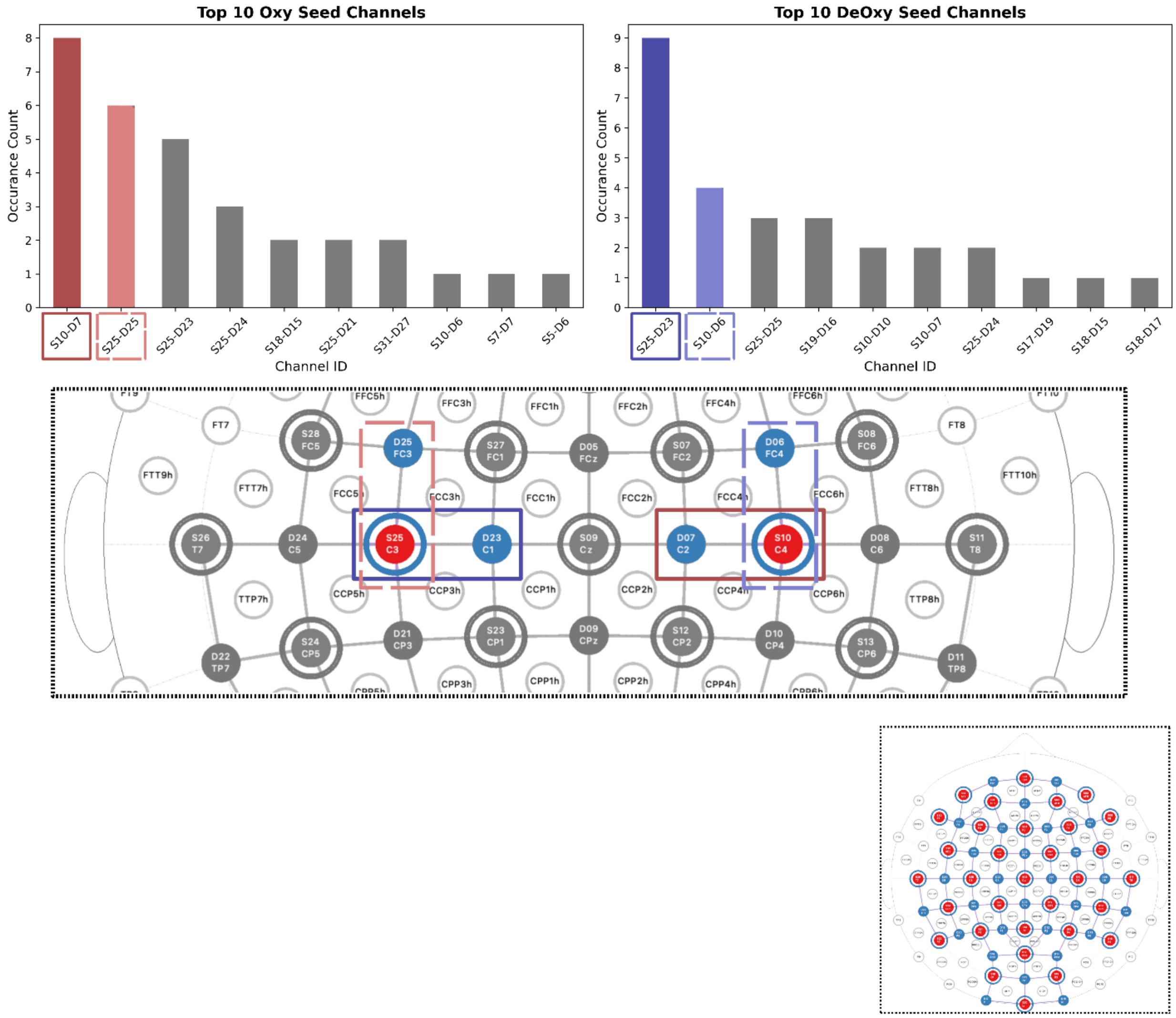


**Fig. S3** The top row shows histograms of the 10 most frequently selected channels for HbO (left) and HbR (right) across subjects (*N* = 38). The full montage (bottom right) provides a reference for the complete channel layout, illustrating the spatial arrangement of all channels across the scalp. In the cropped montage (middle), the two most representative motor seeds for each signal are highlighted with orthogons: red orthogons indicate HbO seeds, and blue orthogons indicate HbR seeds.

#### *S1.6 Subject-level RSFC map*

In Figure S4 the motor RSFC map from a single subject at random is shown to demonstrate the capability of fNIRS in identifying individualized-RSFC patterns. For this illustration, heatmaps from ICA-LogCosh and SBA-Correlation were selected as an example, while SBA-GLM was not chosen as its performance was significantly lower than the other approaches. Using SBA, the left motor network is clearly identifiable with both chromophores, whereas the right motor network is less distinct. In contrast, ICA allows for a clearer identification of the bilateral motor connectivity in both HbO and HbR. However, ICA also reveals an additional frontal component with asynchronous activity with the motor network. This component is spatially prominent and comparable in magnitude to the motor network, resulting in a more complex and less visually isolated pattern of motor RSFC. This reflects ICA's sensitivity to multiple concurrent sources of connectivity, but at the same time complicates the interpretation of single-network RSFC connectivity maps at the subject level.

While these observations are based on visual inspection and should therefore be interpreted with caution, they showcase fNIRS' potential for single-subject RSFC detection. Future work should aim to systematically evaluate whether individualized-RSFC patterns detected with fNIRS have sufficient spatial specificity to be clinically informative. Ideally, this should be validated using simultaneous fNIRS-fMRI measurements, enabling direct assessment of how closely fNIRS connectivity approximates that of fMRI.

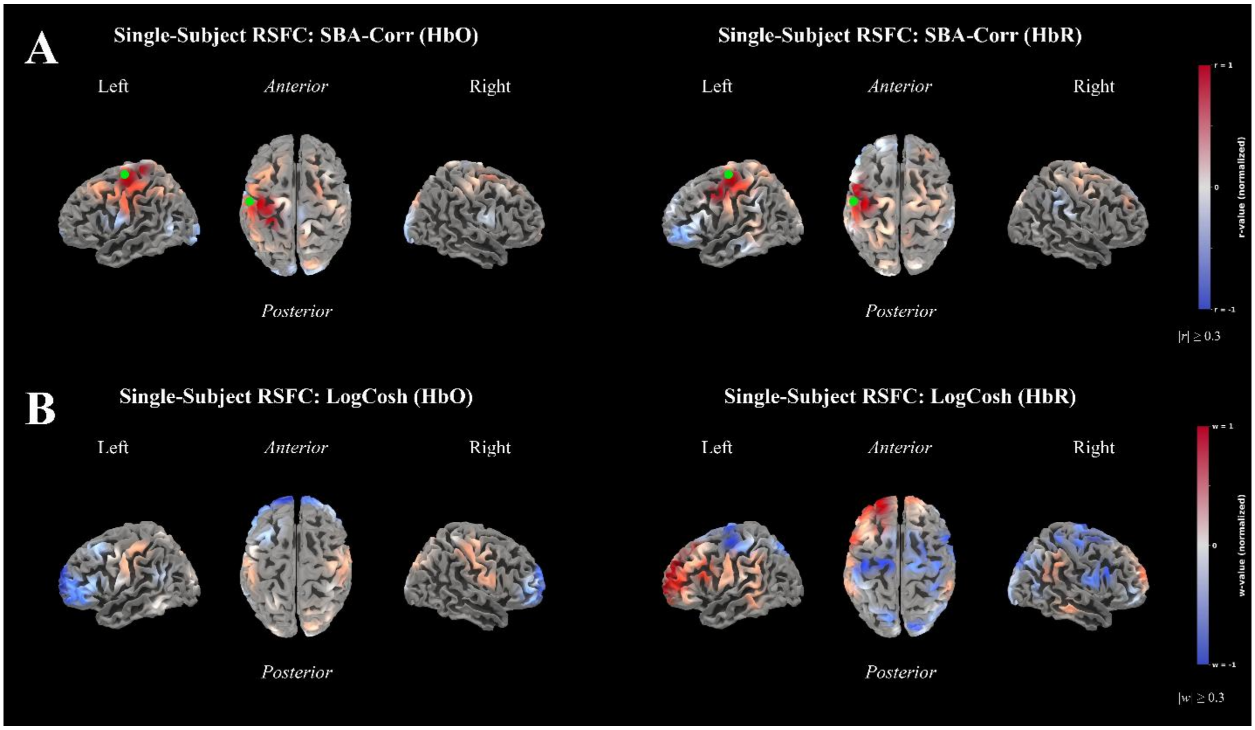


**Fig. S4** (a) SBA-Corr and (b) ICA-LogCosh RSFC heatmaps of HbO and HbR from a single subject at random. For SBA, the seed channel (S25-D23) is illustrated with a green dot on the left hemisphere. For visualization purposes, the *r*-value of the seed channel was capped at the second highest (to avoid visualization distortions), and then all *r*-values were divided by the absolute maximum *r* to produce a normalized map ranging from -1 to +1, comparable to ICA. For ICA, the subject's RSFC map was obtained by taking the median across 50 ICA runs, and then the channel weights were linearly rescaled from -1 to +1. For both SBA and ICA, the values below 0.3 were masked.
